# *ESR1* mutant breast cancers show elevated basal cytokeratins and immune activation

**DOI:** 10.1101/2020.12.29.424777

**Authors:** Zheqi Li, Yang Wu, Amir Bahreini, Nolan M. Priedigkeit, Kai Ding, Carol A. Sartorius, Lori Miller, Margaret Rosenzweig, Nikhil Wagle, Jennifer K. Richer, William J. Muller, Laki Buluwela, Simak Ali, Yusi Fang, Li Zhu, George C. Tseng, Jason Gertz, Jennifer M. Atkinson, Adrian V. Lee, Steffi Oesterreich

## Abstract

Estrogen receptor alpha (ER/*ESR1*) is mutated in 30-40% of endocrine resistant ER-positive (ER+) breast cancer. *ESR1* mutations cause ligand-independent growth and increased metastasis *in vivo* and *in vitro*. Despite the distinct clinical features and changes in therapeutic response associated with *ESR1* mutations, there are no data about their potential role in intrinsic subtype switching. Applying four luminal and basal gene set pairs, *ESR1* mutant cell models and clinical samples showed a significant enrichment of basal subtype markers. Among them, the six basal cytokeratins (BCKs) were the most enriched genes. Induction of BCKs was independent of ER binding and instead associated with chromatin reprogramming centered around a progesterone receptor-orchestrated topological associated domain at the *KRT14/16/17* genomic region. Unexpectedly, high *BCK* expression in ER+ primary breast cancer is associated with good prognosis, and these tumors show enriched activation of a number of immune pathways, a distinctive feature shared with *ESR1* mutant tumors. S100A8 and S100A9 were among the most highly induced immune mediators shared between high-*BCK*s ER+ and *ESR1* mutant tumors, and single-cell RNA-seq analysis inferred their involvement in paracrine crosstalk between epithelial and stromal cells. Collectively, these observations demonstrate that *ESR1* mutant tumors gain basal features with induction of basal cytokeratins via epigenetic mechanisms in rare subpopulation of cells. This is associated with increased immune activation, encouraging additional studies of immune therapeutic vulnerabilities in *ESR1* mutant tumors.

## Introduction

Breast cancer is characterized by a high degree of heterogeneity, originally identified through the use of immunohistochemistry and gene expression profiling^1,2^. Broadly, molecular subtypes can be grouped into luminal (luminal A and luminal B), HER2-enriched and basal-like tumors, primarily driven by expression of ER, PR and HER2 and Ki67^3^. Tumors with different molecular subtypes show distinguishing clinical features and therapeutic responses^4,5^, including metastatic spread and immune profiles^6,7^.

The basal-like subtype, which represents 15-25% of all cases and overlaps with triple negative breast cancers (TNBC), is characterized by a unique gene expression profile similar to that of myoepithelial normal mammary cells^8^. Basal-like breast cancers are more aggressive and patients suffer from shorter metastases-free survival compared to those with luminal subtypes^8,9^. Mechanisms underlying increased invasive properties of basal-like tumors include deregulation of the CCL5/CCR5 axis^10^, amplified EGFR^11^ kinase signaling and activation of TGF-β signaling^12^. Despite multiple signaling aberrations providing challenges for efficient therapeutic strategies, recent studies have unveiled unique vulnerabilities of basal-like breast cancers, such as higher levels of PD-L1 expression along with constitutive IFNγ signaling activation^13^, in line with higher immune- infiltration scores^6^. While the FDA has granted an accelerated approval for atezolizumab, a monoclonal antibody drug targeting PD-L1, plus chemotherapy for the treatment of TNBC^14^, the potential application of immune therapies for patients with luminal breast cancer remains largely unknown.

Among the four intrinsic subtypes, basal and luminal subtypes show opposite histochemical features and notable differences in prognosis^15,16^, however there is increasing evidence that these subtypes are on a continuum of “luminal-ness” and “basal-ness” features. Models of breast cancer lineage evolution describe that basal and luminal progenitor cells are derived from the same bipotential progenitors^17^, indicating the potential of lineage reprogramming during cancer progression. Such subtype switching during tumor evolution has been described and is critical for implementation of precision therapeutics^18–20^. A recent study by Bi et al. reported loss of luminal and gain of basal markers in endocrine resistant breast tumors^21^. Mechanisms underlying the intrinsic subtype plasticity are largely unknown, with some exceptions. *JARID1B*^22^ and *ARID1A^23^* have been described as essential luminal lineage driver genes and their mutations result in luminal-to-basal subtypes switches. In addition, enhancer reprogramming at GATA3 and AP1 binding sites has been highlighted as a pivotal epigenetic mechanism allowing lineage plasticity^21^.

ER is well characterized as a luminal lineage marker^24^. Hotspot mutations in its ligand-binding domain occur in 30%-40% of endocrine resistant breast tumors, promoting ligand-independent ER activation and metastasis^25–27^. Several recent studies showed that *ESR1* mutant tumors are not only associated with endocrine resistance, but also gain unexpected resistance towards CDK4/6 inhibitors^28^, mTOR inhibitors^29^ and radiation therapy^30^ in a mutation subtype and context dependent manner, suggesting potentially more complex re-wiring of ER mutant tumors.

We set out to examine whether *ESR1* mutations alter the “luminal-ness” and “basal-ness” balance in breast cancer cell line models and clinical specimens. We discovered that ER mutant tumors gain basal-like features, characterized by elevated expression of basal cytokeratins as a result of epigenetic reprogramming. Immune context analyses in clinical specimens revealed potential therapeutic vulnerabilities accompanying the increased basal-ness in *ESR1* mutant breast cancer, a finding of potential clinical relevance.

## Results

### Basal gene signatures are enriched in *ESR1* mutant breast cancer

To examine whether *ESR1* mutations alter “luminal-ness” and “basal-ness” we utilized four independent luminal and basal gene signatures (Fig. 1A, Supplementary Table S1). Gene sets from Charafe-Jauffret et al.^31^ and Huper et al.^32^ were obtained from MSigDB (Supplementary Fig. S1A and S1B), and in addition we generated two other gene sets from i) intrinsic subtype genes^33^ differentially expressed between luminal (n=33) and basal (n=39) breast cancer cell lines (Supplementary Table S2) ^34–36^ and ii) genes differentially expressed between luminal and basal primary tumors in TCGA ^37^ (Supplementary Fig. S1C and S1D). Although the overlap among the different gene sets was limited (Fig.1B), likely reflecting differences in methodology and sources, some well described lineage marker genes (e.g. *ESR1* and *FOXA1* as luminal markers, and *KRT6A* and *KRT16* as basal markers) were observed in 3 out of 4 gene sets.

**Figure 1.**
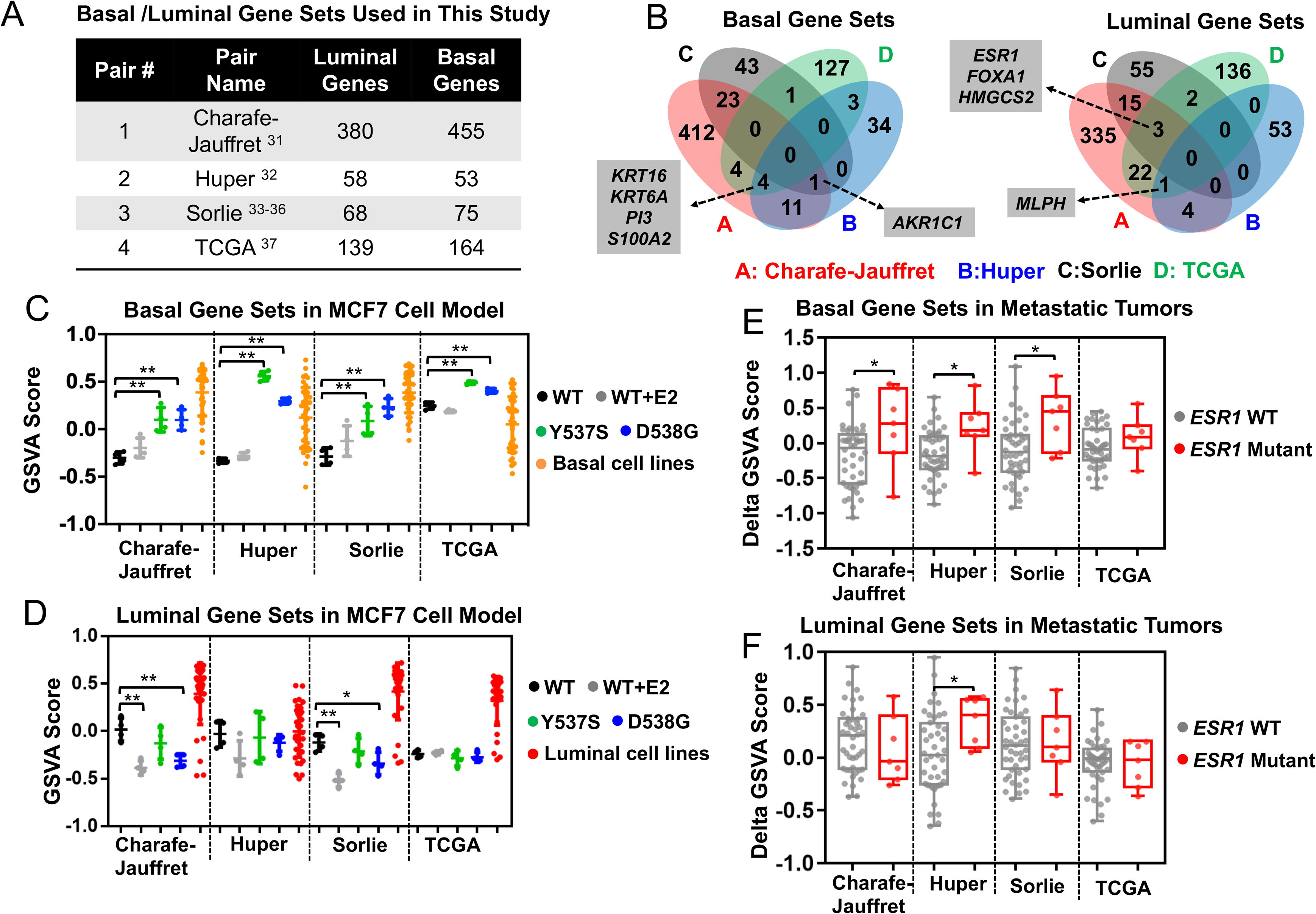
Basal breast cancer gene sets are enriched in *ESR1* mutant breast cancers. A) Four pairs of luminal/basal gene sets applied in this study with gene numbers specified in each set. B) Venn diagram representing the overlap of genes from the basal (left) and luminal (right) gene sets. Genes overlapping in at least three gene sets are indicated. C) and D) Dot plots showing GSVA score of the four pairs of basal (C) and luminal (D) gene sets enrichment in MCF7 genome-edited cell models. Scores from luminal and basal breast cancer cell lines were used as positive controls. Dunnett’s test was used to compare with WT-vehicle set within each gene set. (* p<0.05, ** p<0.01) E) and F) Box plots representing basal (E) and luminal (F) gene set enrichments in intra-patient matched paired primary-metastatic samples. Delta GSVA score for each sample was calculated by subtracting the scores of primary tumors from the matched metastatic tumors. Mann-Whitney U test was performed to compare the Delta GSVA scores between WT (N=44) or *ESR1* mutation-harboring (N=7) paired tumors. (* p<0.05)

As expected, all four basal gene sets were significantly enriched in basal versus luminal breast cancer cell lines and tumors (Supplementary Fig. S2A and S2B), and *vice versa* for luminal gene sets except for the Huper luminal markers, likely due to its derivation from normal mammary tissue (Supplementary Fig. S2C and S2D). We found concordantly increased enrichment of basal gene sets in Y537S and D538G MCF7 *ESR1* genome-edited mutant cells, whereas no differences were observed in estrogen treated *ESR1* wildtype cells (Fig. 1C). In contrast, we did not observe a consistent change in the luminal gene sets (Fig. 1D). The enrichment of the basal gene sets in the *ESR1* mutant cells was also seen in an independent CRISPR-engineered MCF7 *ESR1* mutant cell model recently reported by Arnesen et al^38^ (Supplementary Fig. S3A) and in our T47D *ESR1* mutant cells^27^ (Supplementary Fig. S3B). Of note, no consistent and strong alterations of luminal and basal gene sets enrichment levels were detected in *ESR1* WT endocrine resistant ER+ breast cancer cell models ^21,39–46^ (8 tamoxifen resistant, 2 fulvestrant resistant and 7 long-term estradiol deprivation (LTED) models), suggesting that the “basal-ness” shift is a unique feature acquired as a result of *ESR1* mutations (Supplementary Fig. S3C)^46^.

We next sought to extend our findings to clinical specimens using RNA-seq data composed of 51 intra-patient matched ER+ primary-metastatic tumor pairs (7 *ESR1* mutant and 44 *ESR1* WT pairs) (Supplementary Table S3). Similar to observations in cell lines, *ESR1* mutant metastatic breast cancers showed a significant enrichment of basal gene signatures compared to tumors with WT *ESR1* (Fig. 1E). We did not observe a concurrent decrease of luminal markers (Figure 1F). Taken together, these findings suggested a novel and unexpected gain of “basal-ness” in *ESR1* mutant tumors.

### Basal cytokeratins are elevated in *ESR1* mutant breast cancer cells and tumors

We next interrogated the union of the four basal gene sets (N=634) to identify which basal marker genes were consistently induced in *ESR1* mutant breast cancer cells. Integrating RNA-seq results from MCF7 cell models^27^ and clinical samples identified a group of basal cytokeratins (*KRT5, KRT6A, KRT6B, KRT14, KRT16, and KRT17*) as the top consistently increased basal markers (Fig. 2A, Supplementary Fig. S4A and Supplementary Table S4). Elevated basal cytokeratins (BCKs) mRNA levels were further confirmed in independent qRT-PCR experiment in *ESR1* mutant MCF7 cells (Fig. 2B). Analyzing fold-change expression of all basal markers in a number of MCF7 *ESR1* mutant cell models previously described^25,27,38^ revealed *KRT5,16* and *17* as the top increased basal genes (Supplementary Fig. S4B-D). In the T47D *ESR1* mutant cells, *KRT16* was significantly increased (Supplementary Fig. S4E), but the observed enrichment of basal marker genes (Supplementary Fig. S3B) was also driven by other non-canonical basal genes such as *WLS and HTRA1* (Supplementary Table S5), suggesting some context-dependent mechanisms for the increased basal-ness.

**Figure 2.**
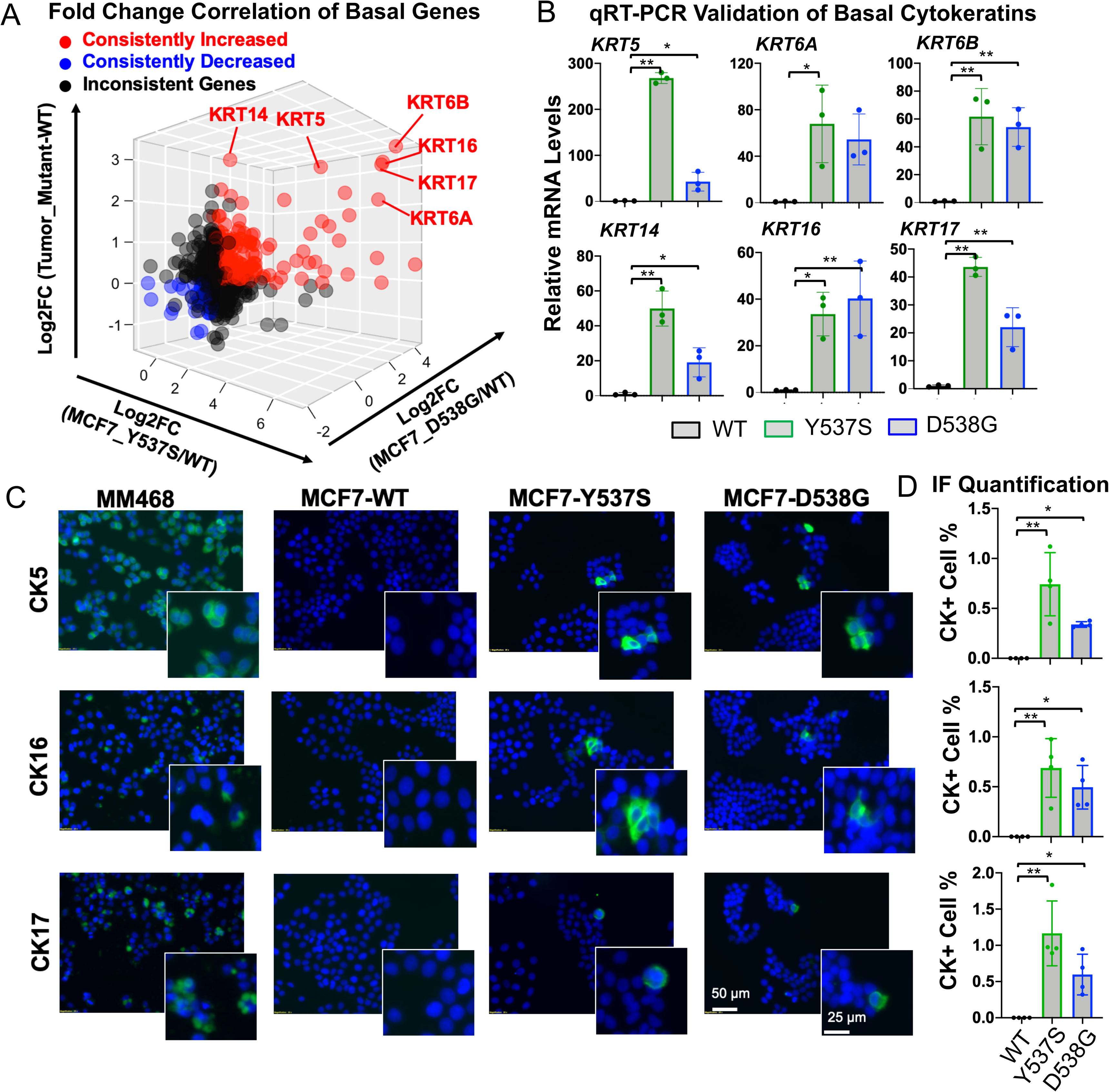
Overexpression of basal cytokeratins (BCK) in *ESR1* mutant breast cancer cells and tumors. A) Correlation between basal gene fold changes (FC) in MCF7-Y537S/D538G cells (normalized to WT vehicle) and intra-patient paired mutant tumors (normalized to WT tumors) (N=634). Consistently increased or decreased genes in the two MCF7 mutant cells and tumors compared to their WT counterparts were highlighted in red or blue respectively, and six basal cytokeratin genes are indicated. Inconsistently changed genes among the three comparisons are labelled in black. B) *KRT5/6A/6B/14/16/17* mRNA levels in MCF7 WT and *ESR1* mutant cells. Relative mRNA fold change normalized to WT cells and *RPLP0* levels measured as the internal control. Each bar represents mean ± SD with three biological replicates. Representative results from three independent experiments are shown. Dunnett’s test was used to compare BCKs expression levels between WT and mutant cells. C) Representative images of immunofluorescence staining on CK5, CK16 and CK17 in MCF7 WT and *ESR1* mutant cells. Regions with CK positive cells were highlighted in the magnified images. MDA-MB-468 was included as positive control. Images were taken under 20x magnification. D) Quantification of percentages of CK positive cells in MCF7 WT and *ESR1* mutant cells. Each bar represents mean ± SD from four different regions. Data shown are from one representative experiment of three independent experiments. Dunnett’s test was used to compare BCKs positive cell prevalence between WT and mutant cells. (* p<0.05, ** p<0.01)

We also queried *KRT* expression in overexpression models. In MCF7 cells with stable overexpression of HA-tagged WT and mutant ER (Y537S and D538G) (Supplementary Figure S5A and S5B), we again observed significant overexpression of *KRT5, KRT6A, KRT6B, KRT16*, and *KRT17* (Supplementary Fig. S5C).

Given higher BCK mRNA expression in *ESR1* mutant cells, we examined their expression at the protein level. We confirmed higher CK5 and CK16 protein levels in early passage (P6-8) *ESR1* mutant cells, but curiously expression was not detectable in later passages (P30-32) (Supplementary Fig. S6A). This finding was consistent with prior reports on slower growth of CK5+ sub-populations^47^, reflecting selection forces eliminating BCK-positive subclones from luminal cell populations. To determine whether BCK expression was limited to minor sub-populations in *ESR1* mutant cells, we performed IF staining for CK5, CK16 and CK17 in early passage cells (below P12) (Fig. 2C). No BCK positive clones were identified in MCF7-WT cells, while 0.5-1% of Y537S and D538G *ESR1* mutant cells exhibited strong diffuse cytoplasmic CK5/16/17 expression. In addition, 3-5% of *ESR1* mutant cells displayed strong BCK signals localized as foci adjacent to the nucleus (Supplementary Fig. S6B), and this was again not observed in the WT cells. Furthermore, co-staining of CK5+CK16 and CK16+CK17 showed that the BCK proteins were predominantly (in 75%-90% imaged cells) upregulated in the same sub-population of cells (Supplementary Fig. S6C and S6D). In contrast, luminal cytokeratin CK8 was homogenously expressed with stronger expression at the edges of each cell cluster (Supplementary Fig. S6E), suggesting that the marked heterogeneity was a unique feature for BCK expression in the luminal cell background.

### BCK induction is independent of mutant ER DNA binding but requires low ER expression

Mutant ER can function in a ligand-independent manner ^26,27^, and we thus tested whether induction of BCKs resulted from ligand-independent ER activity. We interrogated eight publicly available RNA-seq and microarray data sets with estradiol (E2) treatment in six different ER+ breast cancer cell lines ^26,27,48–51^. In contrast to strong E2 induction of classical ER target genes such as *GREB1, TFF1* and *PGR*, expression of basal and luminal cytokeratins genes was not regulated by E2 with the exception of *KRT7* (Fig. 3A). We then examined whether BCK expression was regulated via *de novo* genomic binding of mutant ER at BCK genes. We performed ChIP-seq in MCF7 WT and *ESR1* mutant cells in the absence and presence of E2. As expected, in the absence of E2 we detected very few ER binding sites in WT MCF7 cells (n=125), whereas E2 stimulation triggered substantial ER binding events (n=12,472) (Supplementary Table S6). Consistent with previous studies^25,26^, Y537S and D538G ER show strong ligand-independent binding, with 657 binding sites in Y537S and 1,016 in D538G mutant cells (Supplementary Fig. S7A). The *GREB1* gene locus is shown as a representative example (Fig 3B, left panel). Co-occupancy analyses between WT-E2 and mutant-vehicle sets demonstrated that one third of all Y537S (36%) and D538G (31%) ER binding sites were not detected in the WT+E2 data suggesting gain-of-function novel binding sites (Supplementary Fig. S7B); however, none of them mapped to the BCKs genes with increased expression in *ESR1* mutant cells (−/+ 50kb of transcriptional start sites) (Fig. 3B, middle and right panel).

**Figure 3.**
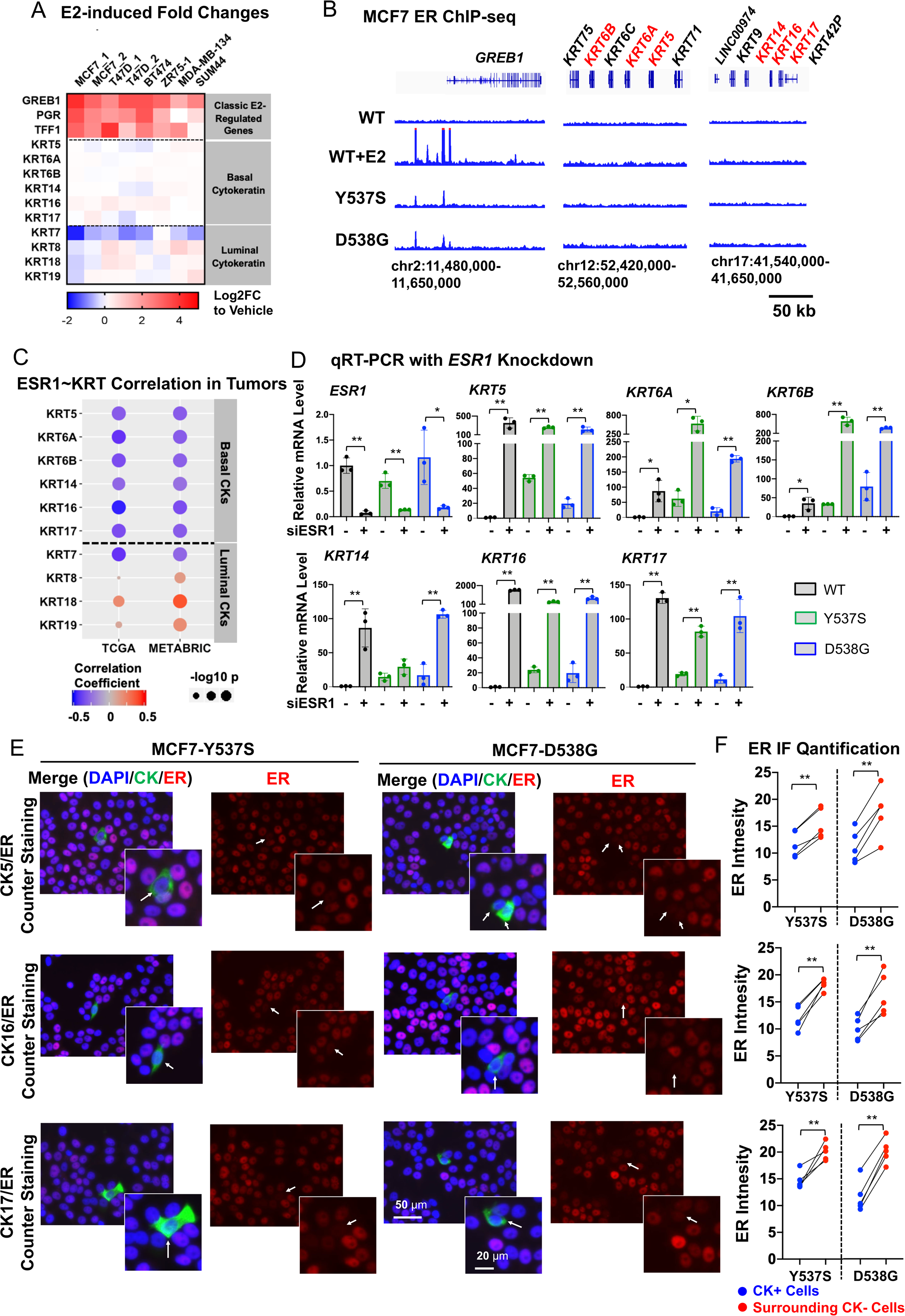
Basal cytokeratins induction is independent of mutant ER genomic binding but requires low ER expression. A) Heatmap representing fold change mRNA expression (E2/veh) of six basal cytokeratins and four luminal cytokeratins in ER+ breast cancer lines from six publicly available data sets (GSE89888, GSE94493, GSE108304, GSE3834, GSE38132 and GSE50693). *GREB1*, *PGR*, and *TFF1* are canonical E2-regulated genes included as positive controls. B) Genomic track showing ER binding intensities at *KRT5/6A/6B* and *KRT14/16/17* loci from ER ChIP-seq data sets of MCF7 *ESR1* mutant cells. *GREB1* locus serve as a positive control. C) Graphic view of Pearson correlation between expression of *ESR1* and each basal or luminal cytokeratin in ER+ breast tumors in TCGA (n=808) and METABRIC (n=1,505) cohorts. Color scale and size of dots represent correlation coefficient and significance, respectively. D) qRT-PCR measurement of *ESR1*, *KRT5/6A/6B/14/16/17* mRNA levels in MCF7 WT and *ESR1* mutant cells with *ESR1*siRNA knockdown for 7 days. mRNA fold change normalized to WT cells; RPLP0 levels were measured as internal control. Each bar represents mean ± SD with three biological replicates. Data shown are representative from three independent experiments. Student’s t-test was used to compare the gene expression between scramble and knockdown groups. (* p<0.05, ** p<0.01) E) Representative images of ER, CK5, CK16 and CK17 staining in MCF7-Y537S and D538G cells. BCKs positive cells are highlighted with white arrows. Images were taken under 20x magnification. F) Bar plots quantifying the ER intensities in BCKs positive (blue) and the corresponding proximal negative (red) cells from each region. Each bar represents mean ± SD analyzed in five different regions per group from one experiment, representative of three independent experiments. Paired t test was applied to compare ER intensities between BCKs positive and negative cells. (* p<0.05, ** p<0.01)

We then expanded our analyses and examined potential estrogen-regulation of all basal marker genes, again using the union of the four basal gene sets (N=634). Comparison of E2 and *ESR1* mutation-conferred fold changes of these genes in MCF7 cells revealed that the top upregulated basal markers in *ESR1* mutant cells were not E2-induced (Supplementary Fig. S7C and S7D). In addition, only 20 basal genes (3%) harbor mutant ER binding sites at −/+ 50 kb of TSS (Supplementary Fig. S7E), and 18 of those were not differentially expressed between WT and mutant cells (Supplementary Fig. S7F). Taken together, these analyses suggest that the shift to “basal-ness” in *ESR1* mutant cells was not mediated via ligand-independent binding of mutant ER to BCK gene loci.

To further understand interplay between *ESR1* and *KRT* gene expression, we determined expression of basal and luminal *KRT* genes in ER+ primary breast tumors. As shown in Figure 3C, the six BCKs were significantly negatively correlated with *ESR1* expression, whereas the luminal *KRT* were mostly positively correlated with *ESR1* (Fig. 3C). Luminal *KRT7* was again the exception, being negatively correlated with *ESR1* expression, in line with it being repressed by ER (Figure 3A). The inverse correlation between BCK and *ESR1* expression was also reflected in results from ER knockdown experiments, in which loss of *ESR1* significantly increased expression of BCKs in MCF7 WT and mutant cells (Fig. 3D). Similar results were obtained in five additional ER+ breast cancer cell lines where we observed a general increase of BCK expression after *ESR1* knockdown (Supplementary Fig. S8). In addition, co-staining of ER and CK5/CK16/17 in MCF7 *ESR1* mutant cells showed significantly lower ER expression in BCK+ cells than in the surrounding BCK- cells (Fig. 3E). Collectively, these data demonstrate that ER serves as a negative regulator of BCKs expression independent of ligand and mutational status, and suggest that low ER expression is likely necessary but not sufficient to facilitate BCKs overexpression in a subpopulation of *ESR1* mutant cells. These data also support a role for mutant ER in regulating BCK expression via epigenetic regulation, a mechanism that we have recently shown to be used by mutant ER ^38^.

### PR regulation of BCK expression through binding at a CTCF-driven chromatin loop at the *KRT14*/16/17 loci in *ESR1* mutant cells

To investigate potential epigenetic regulation of *KRT5/6A/6B* and *KRT14/16/17*, we first compared their regional epigenetic landscapes on chromosome 12 and 17, respectively, in luminal and basal breast cancer cell lines and tumors (Supplementary Fig. S9). Integrative analysis of ATAC-seq and ChIP-seq profiles of H3K4me2, H3K4me3, H3K9ac and H3K27ac suggested that these two regions are epigenetically silent in MCF7 (Supplementary Fig. S9A), consistent with low expression. In basal breast cancer cell lines and tumors, there is an enrichment of H3K27 acetylation (Supplementary Fig. S9B) and number of ATAC-seq peaks (Supplementary Fig. S9C) at BCK loci, consistent with increased mRNA expression (Supplementary Fig. S9E and S9F). This is also observed in *ESR1* mutant cell models (Supplementary Fig. S9G).

We recently reported CCCTC-binding factor (CTCF) motif as one of the top enriched motifs in unique *ESR1* mutant-regulated accessible genomic regions^38^. To determine whether CTCF has a role in the epigenetic regulation of BCK, we developed a CTCF gene signature by identification of the top 100 differentially expressed genes before and after CTCF knockdown in MCF7^52^ (Supplementary Table S1). The positively correlated CTCF signature (i.e. using genes that were repressed after CTCF knockdown) was significantly enriched in both MCF7 *ESR1* mutant cells (Fig. 4A) and metastatic tumors (Fig. 4B) compared to their WT counterparts, whereas E2 stimulation had no effect (Fig. 4A). CTCF is a multimodal epigenetic regulator in breast cancer^53^, in part through generating boundaries of topological associating domains (TADs) and guiding of DNA self-interaction^54^. Mapping the genomic occupancy of CTCF and three other cohesion complex members (RAD21, STAG1 and SMC1A) in MCF7 cells^55–57^ (Fig. 4C) identified five putative TAD boundaries at the *KRT14/16/17* (Fig. 4D) loci and three at the *KRT5/6A/6B* (Supplementary Fig. S10A) loci. Integration of an additional MCF7 CTCF ChIA-PET dataset^58^ showed that a strong chromatin loop is predicted to span the *KRT14/16/17* genes, further supported by the pattern of convergent CTCF motif orientations at the predicted TAD boundaries (Fig. 4C). Since the *KRT5/6A/6B* locus did not harbor strong chromatin loops (>3 linkages), we focused our further analysis on the *KRT14/16/17* locus.

**Figure 4.**
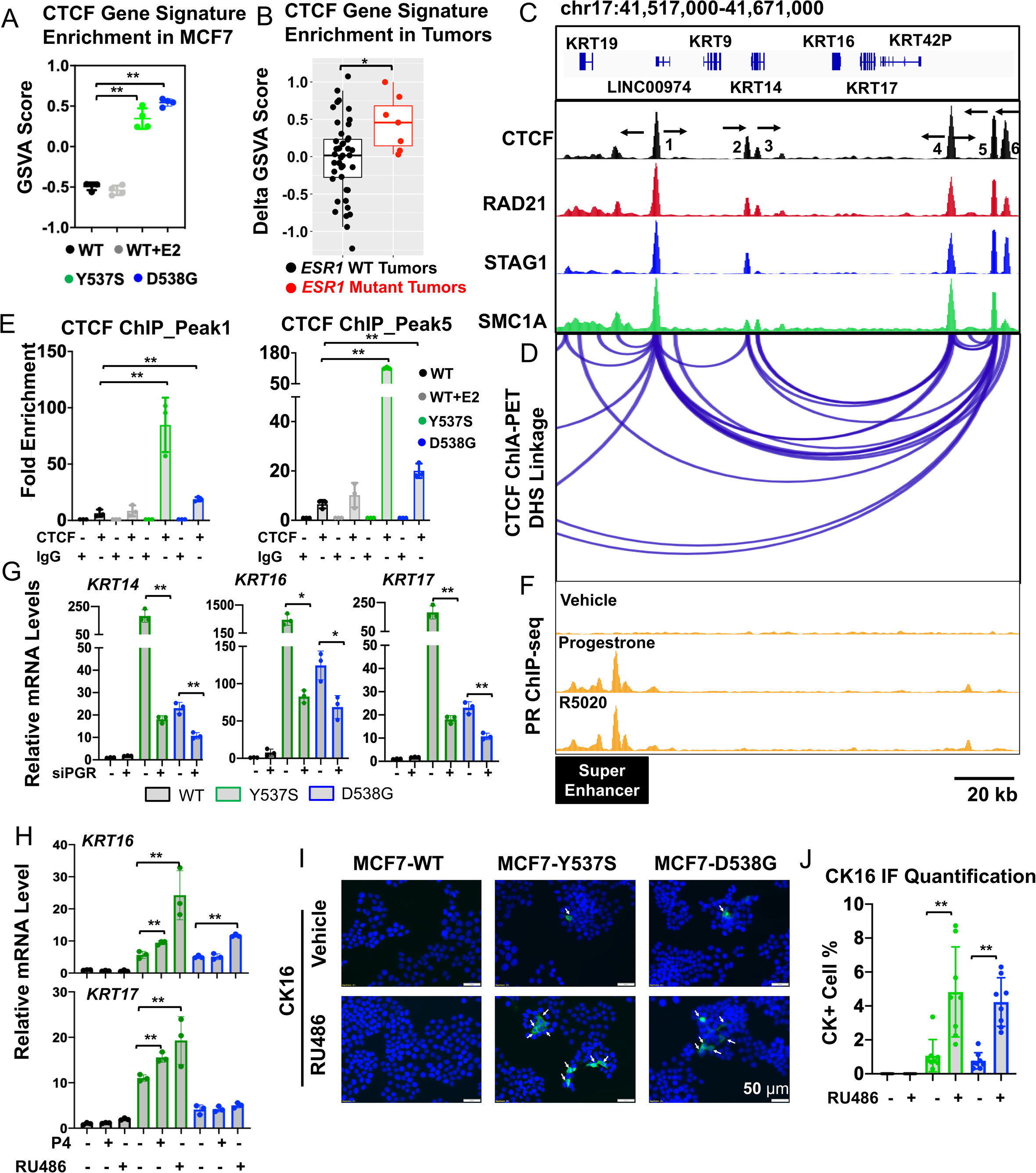
Basal cytokeratins are induced via a unique PR enhancer-associated TAD in *ESR1* mutant cells. A) Dot plots showing enrichment levels of CTCF gene signature in MCF7 *ESR1* mutant cells. Dunnett’s test was used to compare the difference. (** p<0.01) B) Dot plots showing enrichment levels of CTCF gene signature in *ESR1* WT (n=44) and mutant (n=7) metastases. Mann-Whitney U test was used to compare enrichment levels in tumors. (* p<0.05) C) Genomic track illustrating the CTCF/cohesion complex binding at *KRT14*/16/17 proximal genomic region in MCF7 cells. CTCF and RAD21 ChIP-seq were downloaded from ENCODE (ENCSR560BUE and ENCSR703TNG). STAG1 and SMC1A ChIP-seq data were from GEO (GSE25021 and GSE76893). CTCF motif orientations of each peak is labelled with black arrows in the CTCF track. Y-axis represents signal intensity of each track. D) CTCF-driven chromatin loops visualized using a CTCF ChIA-PET data set in MCF7 cells (GSE72816) at the 3D Genome Browser platform. Each linkage represents a chromatin loop. E) Bar graphs displaying CTCF binding events measured by ChIP-qPCR at binding sites 1 and 5 illustrated in (C). CTCF binding fold enrichments were normalized to the average of IgG binding. Each bar represents mean ± SD of fold changes from three independent experiments. Pair-wise t-test on CTCF binding fold enrichment between WT and each mutant was performed. (* p<0.05, ** p<0.01) F) PR binding under R5020 and progesterone treatments visualized based on a reported PR ChIP-seq data set in MCF7 cells (GSE68359). Y-axis represents signal intensity of each track and is adjusted to the same scale. Super enhancer range was highlighted below the genomic track. G) qRT-PCR measurement of *KRT14*, *16* and *17* mRNA levels in MCF7 *ESR1* WT and mutant cells with PGR siRNA knockdown for 7 days. mRNA fold change normalized to WT cells; RPLP0 levels were measured as internal control. Each bar represents mean ± SD with three biological replicates. Data shown are representative from three independent experiments. Student’s t-test was used to compare the gene expression between scramble and knockdown groups. (* p<0.05, ** p<0.01) H) qRT-PCR measurement of *KRT5*, *16* and *17* mRNA levels in MCF7 *ESR1* WT and mutant cells treated with 0.1% EtOH (vehicle),100 nM P4 or 1 μM RU486 treatment for 3 days. mRNA fold change normalized to WT cells; RPLP0 levels were measured as internal control. Each bar represents mean ± SD with three biological replicates. Data shown are representative from three independent experiments. (* p<0.05, ** p<0.01) I) Representative images of immunofluorescence staining of CK5 and CK16 in MCF7 WT and *ESR1* mutant cells after 3 day treatment with 1% EthOH (vehicle) or 1 μM RU486. Images were taken under 20x magnification. J) Quantification of the percentages of CK positive cells in MCF7 cells. Each bar represents mean ± SD from eight different regions combining from two independent experiments. Student’s t test was used to compare % BCK+ cells before and after treatment. (* p<0.05, ** p<0.01)

ChIP revealed strong enrichment of CTCF binding at the base of the chromatin loops of the *KRT14/16/17* locus in *ESR1* mutant cells, however there was a lack of E2 regulation (Fig. 4E). Decreasing CTCF levels led to increased expression of *KRT14, KRT16* and *KRT17* mRNA levels in *ESR1* mutant cells, potentially reflecting a role for CTCF as “classical” insulator, suppressing high expression of these BCKs through the identified super enhancer at the *KRT14, KRT16* locus (Figure 4F). Given identification of progesterone receptor (PR) binding sites within this super enhancer, PR’s previously identified role in regulating *KRT5* expression in luminal breast cancer cells^47,59^, and finally its overexpression in multiple *ESR1* mutant cell models ^25–27,60^ (Supplementary Fig. S10C and S10D), we tested whether PR regulates *KRT14/16/17* expression.

PR ChIP-seq revealed a ligand-inducible PR binding sites in MCF7 cells approximately 32kb upstream of the *KRT14*/*16*/*17* loop region^61^ (Fig. 4F). This PR binding site overlapped with a curated super-enhancer in MCF7 cells^62^, which was additionally supported by strong active histone modifications (Fig. S9). Knockdown of PR partially rescued the increased expression of *KRT14*, *16* and *17* in both *ESR1* mutants (Fig. 4G and Supplementary Fig. S10E). We also observed a similar rescue effect for *KRT5* (Supplementary Fig. S10E), consistent with previous studies^59^. Furthermore, both PR agonist (P4) and antagonist (RU486) treatment increased *KRT5*, *16* and *17* expression in Y537S *ESR1* mutant cells, while only RU486 triggered *KRT5* and *KRT16* expression in D538G mutant (Fig. 4H and Supplementary Fig. S10F). The marked induction effect of RU486, a PR antagonist, is likely due to its previously reported partial agonism via recruitment of coactivators^63^. The RU486-induced CK5 and CK16 increase was further examined by IF, where CK5 (Supplementary Fig. S10G) and CK16 (Fig. 4I and 4J) positive cells increased from 1% to 5%. Of note, CK17 positive cells were not increased by RU486 treatment (Supplementary Fig. S10G), suggesting translational efficiency differences between different BCK subtypes. Together, these data demonstrated that elevated PR expression in *ESR1* mutant cells was essential for BCKs induction, and this was possibly due to an orchestration with a super enhancer which is accessible to regulate *KRT14*/*16*/*17* genes via the CTCF-driven chromatin loop.

### Enhanced immune activation, associated with S100A8-S100A9 secretion and signaling in *ESR1* mutant tumors

Finally, we investigated whether the increased expression of basal genes in *ESR1* mutant tumors confers basal-like features and potentially novel therapeutic vulnerabilities. To identify basal cytokeratin-associated pathways enriched in ER mutant tumors, we at first identified ER+ tumors with the top and bottom quantile of BCK gene enrichment and then computed hallmark pathways differentially enriched between these two groups (Supplementary Fig. S11A). Intersection of these BCKs-associated pathways with those enriched in *ESR1* mutant metastases uncovered seven shared molecular functions, the top four of which are all related to immune responses (Fig. 5A, Supplementary Fig. S11B, S11C and Supplementary Table S7). An orthogonal approach - bioinformatic evaluation using ESTIMATE^64^ - confirmed enhanced immune activation in BCK-high vs BCK-low ER+ tumors albeit still lower than in basal tumors (Fig. 5B). In addition, BCK-high tumors displayed higher lymphocyte and leukocyte fractions according to a recent biospecimens report^65^ (Fig. 5C), and higher *PDCD1* mRNA levels (Supplementary Fig. S11D). Intriguingly, patients with BCK-high ER+ tumors experience improved outcomes (Fig. 5D), and although entirely speculative at this point in time, one could hypothesize that this might be due to increased anti-tumor immune activation.

**Figure 5.**
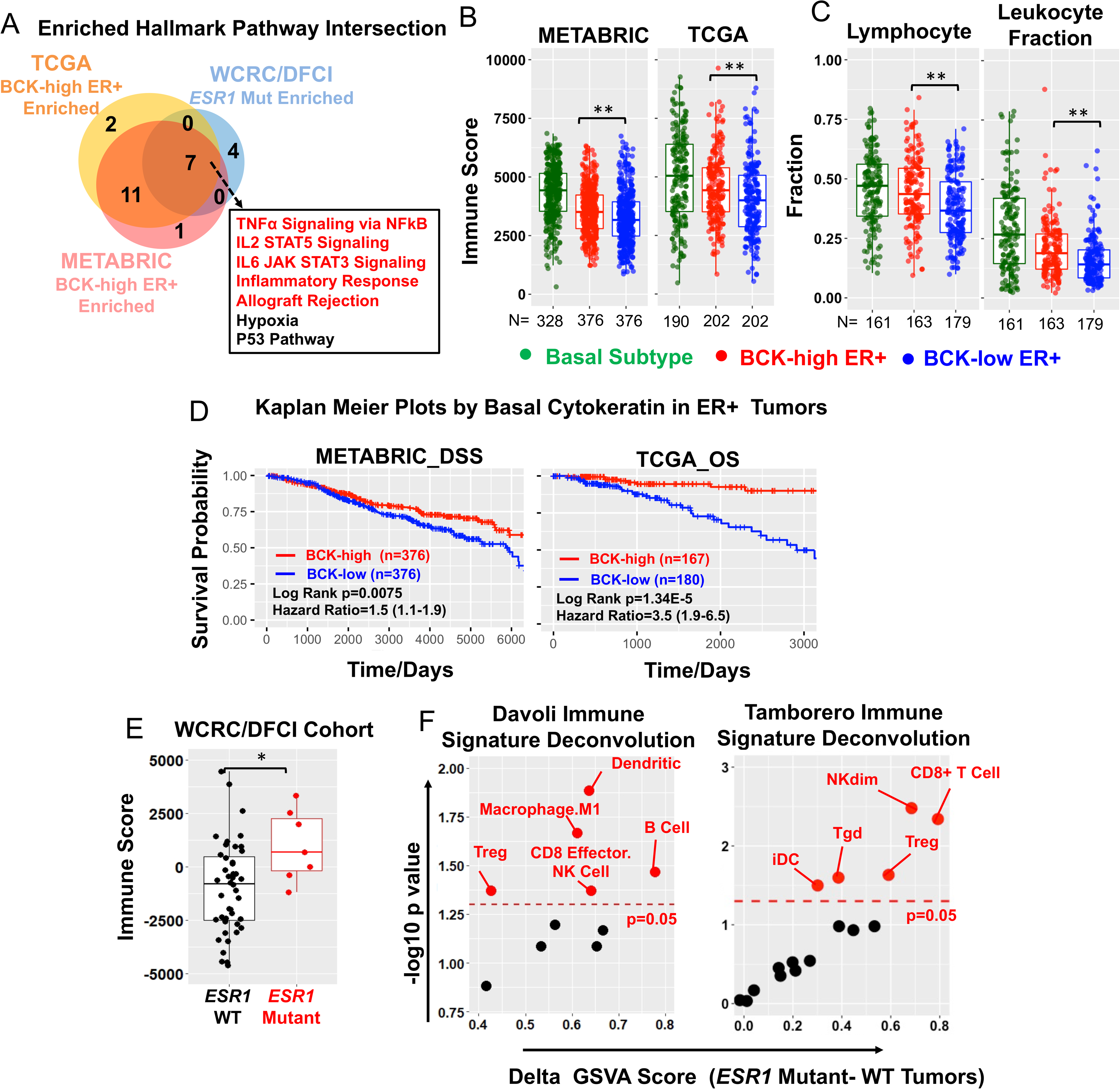
Gain of basal cytokeratin expression is associated with enhanced immune activation in *ESR1* mutant tumors. A) Venn diagrams showing the intersection of significantly enriched hallmark pathways in three sets of comparisons: BCK-high vs low in 1) TCGA ER+ tumors (n=202 in each group), 2) METABRIC ER+ tumors (n=376 in each group) and 3) *ESR1* mutant (n=7) vs WT (n=44) metastatic tumors. BCKs high and low were defined by the upper and bottom quartiles of each subset. The seven overlapping pathways are shown in a frame, and immune-related pathways are highlighted in red. B) Immune scores based on ESTIMATE evaluations in basal tumors (METABRIC n=328; TCGA n=190), BCK-high (METABRIC n=376; TCGA n=202) and low (METABRIC n=376; TCGA n=202) subsets of ER+ tumors in TCGA and METABRIC. Definition of BCK-high and low groups were the same in (A). Mann Whitney U test was used for comparison. (** p<0.01) C) Lymphocytes and leukocyte fractions as determined by a reported TCGA biospecimen dataset^65^ comparing among basal subtype tumors (n=161), TCGA ER+ BCK-high (N=163) and low (N=179) tumors. Definition of BCK-high and low groups were the same in (A). Mann Whitney U test was applied to compare the fractions between BCK-high and low tumors. (** p<0.01) D) Kaplan-Meier plots showing the disease-specific survival (DSS) (METABRIC) and overall survival (OS) (TCGA) comparing patients with ER+ BCKs high vs low tumors. BCKs high and low were defined by the upper and bottom quartiles of each subset. Censored patients were labelled in cross symbols. Log rank test was used and hazard ratio with 95% CI were labelled. E) Immune scores based on ESTIMATE evaluations in *ESR1* mutant (n=7) and WT metastatic (n=44) lesions. Mann Whitney U test was used for comparison. (* p<0.05) F) Dot plot showing the enrichment level alterations of immune cell subtypes in *ESR1* mutant metastatic lesions using Davoli^66^ and Tamborero^67^ immune cell signatures. RNA seq data from intra-patient matched *ESR1* mutant (N=7) and WT (N=44) was used. Immune cell subtypes showing significant increase in *ESR1* mutant tumors were labelled in red (p<0.05).

Similar to BCK-high ER+ tumors, *ESR1* mutant metastatic tumors exhibited higher immune scores compared to those with *ESR1* WT (Fig. 5E). Immune cell subtype deconvolution^66,67^ revealed significantly higher CD8+ T, NK and dendritic cells, along with macrophages in *ESR1* mutant tumors. Basal breast cancers harbor high immune infiltrations at least in part due to higher tumor mutation burden (TMBs)^68^, however, we did not detect higher TMB in BCK-high vs low ER+ tumors (Supplementary Fig. S11E).

To understand which factors might contribute to immune activation in *ESR1* mutant and BCK-high ER+ tumors, we compared gene expression of major immune genes derived from ESTIMATE^69^ (n=141) between *ESR1* mutant and WT tumors, and BCK-high vs BCK-low ER+ tumors. This analysis identified S100A8 and S100A9 as the two top consistently increased immune-related genes (Fig. 6A), and this overexpression was also seen in MCF7 *ESR1* mutant cell models (Supplementary Fig. S11F). S100A8 and S100A9 are pro-inflammatory cytokines that form heterodimers and play crucial roles in shaping immune landscapes^45,46^. As expected, S100A8-A9 expression correlated positively with immune scores in ER+ tumors (Fig. 6B). BCKs levels failed to differentiate immune scores in ER+ tumors among the subset of tumors exhibit high S100A8-A9 (Fig. 6B). S100A8-A9 are secreted proteins and function as heterodimers. To confirm S100A8-A9 protein overexpression, we measured S100A8-A9 heterodimer levels in plasma samples from patients with *ESR1* WT (n=7) and mutant (n=11) tumors (Supplementary Table S8) (Fig. 6C). This analysis revealed significantly higher circulatory S100A8-A9 heterodimers concentrations in plasma from patients with *ESR1* mutations (Fig. 6D).

**Figure 6.**
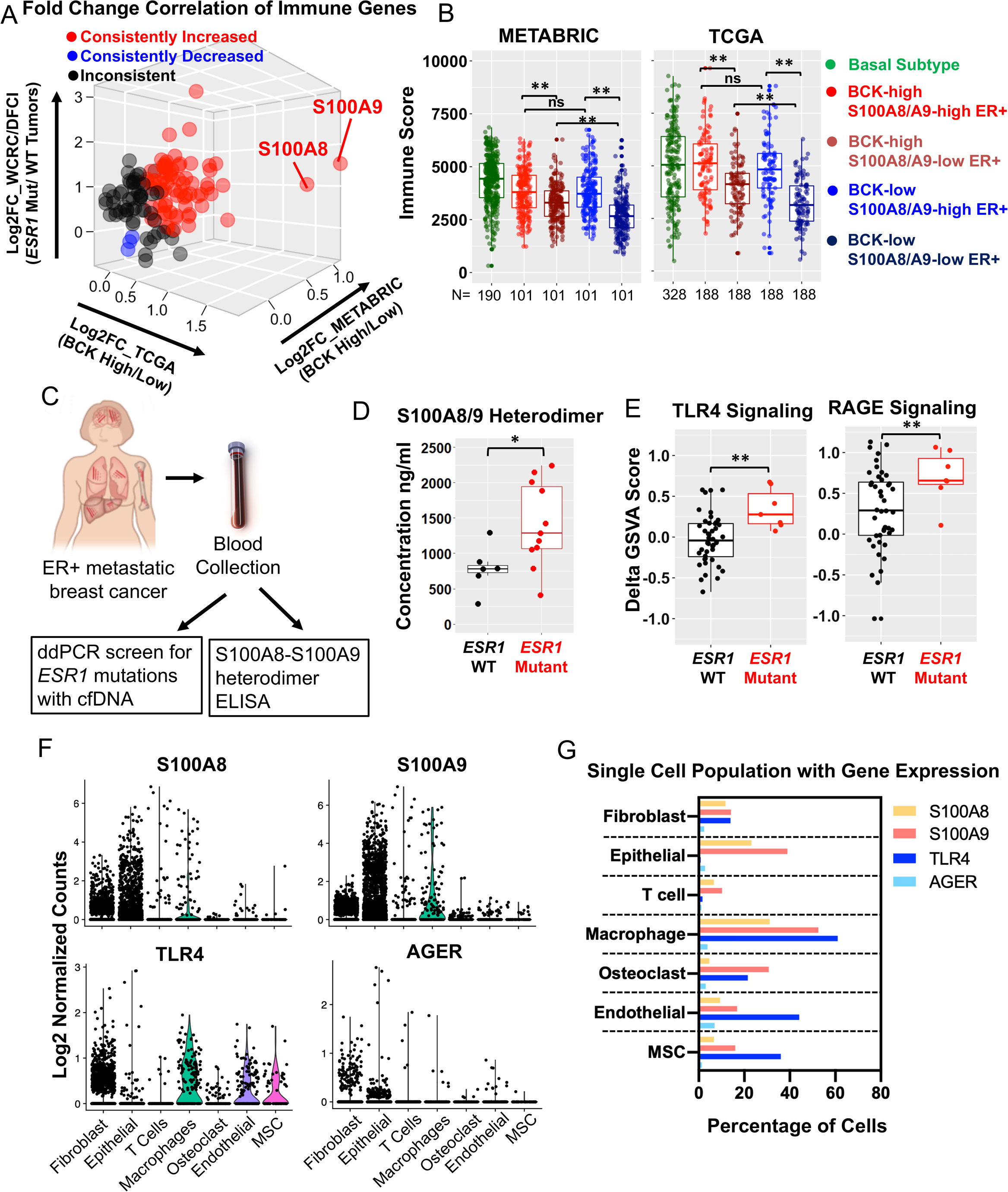
Immune activation in *ESR1* mutant tumors is associated with S100A8/A9-TLR4 paracrine crosstalk between epithelial and stromal cells. A) Three-dimensional plot showing fold change (FC) expression changes of immune genes from ESTIMATE (N=141)^69^ comparing ER+ BCK-high vs low tumors (TCGA and METABRIC) and intra-patient paired *ESR1* WT/mutant tumors. Consistently increased/decreased genes in TCGA and METABRIC BCK-high tumors and *ESR1* mutant tumors were highlighted in red and blue. Inconsistently changed genes among the three comparisons are labelled in black. B) ER+ cases with BCK-high and low quantiles were further divided by the mean expression of S100A8 and S100A9. ESTIMATE immune scores were compared across all four subsets (n=188 and 101 in each group of METABRIC and TCGA) together with basal tumors (n=328 METABRIC and n=190 TCGA). Each corresponding comparison was tested using Mann Whitney U test. (**p<0.01) C) Graphical presentation of strategy to quantify and compare S100A8/9 heterodimer abundance in plasma from patients with ER+ metastatic breast cancer. D) Box plot showing S100A8/9 heterodimer concentrations in plasma from patients with *ESR1* WT (n=7) and mutant (n=11) metastatic breast cancer. Mann Whitney U test was utilized. (* p<0.05) E) Comparison of TLR4 (left) and RAGE (right) signaling signature enrichments in intra-patient matched *ESR1* mutant (N=7) and WT (N=44) cohort. Delta GSVA score of each sample was calculated by subtracting the scores of primary tumors from the matched metastatic tumors. Mann-Whitney U test was performed between WT and mutant tumors. (**p<0.01) F) Violin plots showing S100A8, S100A9, TLR4 and AGER expression by log2 normalized counts in different cell subtypes using single-cell RNA-seq data from two bone metastases from a patient with ER+ breast cancer. G) Percent of cells expressing S100A8, S100A9, TLR4 and AGER, using single cell RNA seq data shown in Figure 6F.

S100A8-A9 heterodimer mainly stimulates downstream cascades through two receptors: toll-like receptor 4 (TLR4) and receptor for advanced glycation end products (RAGE), and both of them are widely reported to impact cancer immunity. A further gene set variation analysis in WCRC/DFCI primary-matched paired metastatic samples revealed consistent enrichment of both pathways in *ESR1* mutant tumors (Fig. 6E, Supplementary Table S1), suggesting both TLR4 and RAGE signaling are hyperactive in *ESR1* mutant tumors.

To further elucidate the specific cell-cell communication by S100A8/S100A9 signaling, we analyzed RAGE and TLR4 signaling via measuring ligand and receptor expression in different cell types using single-cell RNA-seq data from two breast cancer metastases. Highest expression of *S100A8/S100A9* was seen in epithelial cells, followed by fibroblast and macrophages. In contrast, TLR4 and AGER (RAGE) showed low expression in the epithelial cells, but instead were widely expressed in the stroma, especially in fibroblasts and macrophages. In general, AGER displayed lower expression levels in all cell types compared to TLR4 (Fig. 6F and 6G).

Taken together, these data support the concept that the increase in basal-ness of *ESR1* mutant tumors is associated with immune activation, in part facilitated by the paracrine S100A8/A9-TLR4 signaling.

## Discussion

Recurrence of ER+ breast cancer causes over 24,000 deaths each year in the US alone. Given that *ESR1* mutation occur in as many as 20-30% of metastatic recurrences, it is imperative to identify therapeutic vulnerabilities through dissecting mechanisms of action. In this study we have uncovered a previously unrecognized plasticity of *ESR1* mutant cells, reflected by enrichment of basal subtype genes in *ESR1* mutant tumors and in particular a gain of BCK expression, resulting from epigenetic reprogramming of a mutant ER-specific PR-linked chromatin loop. This molecular evolution, i.e. an increase of basal-like feature in the *ESR1* mutant tumors was associated with immune activation including enhanced S100A8/A9-TLR4 signaling (Fig 7).

**Figure 7.**
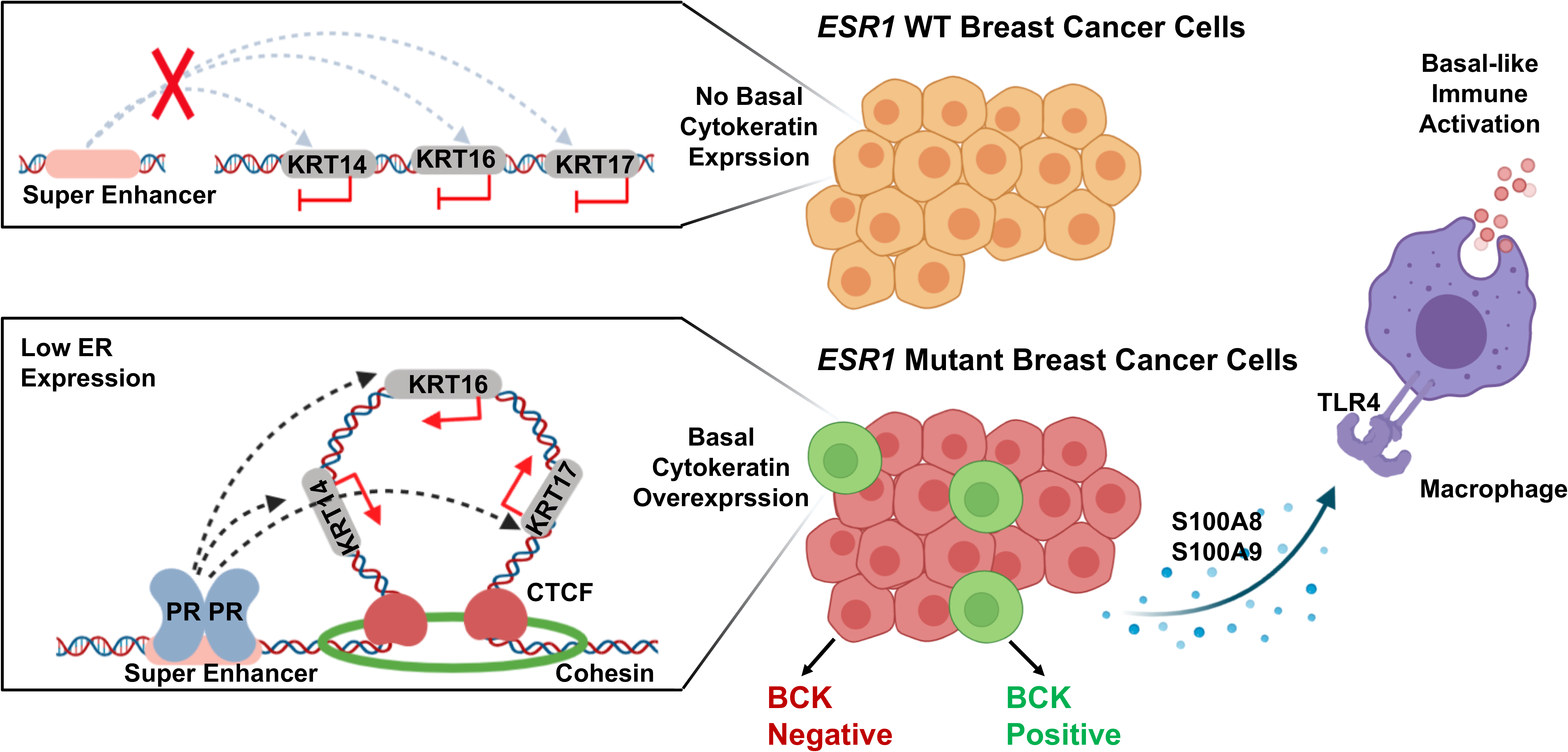
Graphical presentation of proposed mechanisms and relevance of basal cytokeratin induction in *ESR1* mutant breast cancer. *ESR1* WT cells exhibit low basal cytokeratin expression with baseline TAD prevalence spanning *KRT14*/16/17 loci. In contrast, a minor subpopulation of *ESR1* mutant cells exhibit strong basal cytokeratin expression, due to PR activated enhancer at the *KRT14*/16/17 gene locus-spanning TAD. Increased expression of basal cytokeratin is associated with immune activation in *ESR1* mutant tumor similar to that seen in basal tumors, at least in part mediated via enhanced S100A8/A9-TLR4 paracrine crosstalk between epithelial and stromal cells, including macrophages.

Increased plasticity of tumors has previously been shown to be associated with tumor initiation and progression^21,46,70–72^. PAM50 intrinsic subtype switching has been described to occur in as many as 40% of breast cancer metastases^20^. Here we show that *ESR1* mutant cells gain basal-ness, and a similar observation was recently reported by Gu et al.^73^ showing a luminal to basal switch in MCF7 *ESR1* Y537S CRISPR cells compared to parental cells. However, luminal to basal subtype switching is rare in breast cancer^20^ and we have previously reported on clinically relevant gene expression changes in brain metastases (increased in *HER2* gene expression) without clear subtype switching^18^. These results are in line with the increasing appreciation of the molecular subtypes being on a continuum rather than representing discrete stages. Of note, we did not observe a similar gain of basal-ness in a series of *ESR1* wildtype endocrine resistant *in vitro* models, with the exception being a study revealing a “luminal-to-basal” switch in an estradiol-deprived T47D xenograft derived cell line, indicating a potential role for the microenvironment in mediating a similar switch in ER wildtype tumors^74^.

We propose that the observed *ESR1* mutant-cancer cell state interconversions are of potential clinical relevance due to increased stromal immune activation associated with the induction of BCK. Using *in silico* gene expression, pathway analyses and pathology information, we observed increased activation of a number of immune-related pathways including S100A8/S100A9-TLR4 signaling and increased lymphocytic infiltration. S100A8/S100A9 heterodimers exhibit pro-inflammatory properties in different contexts in breast cancer^75,76^, are associated with poor prognosis in multiple cancer types^36^ including breast cancer^77^, and blockade of their activity improves survival^78^. We observed increased S100A8/S100A9 levels in blood from patients with *ESR1* mutant tumors but given complexity of tumor-cell intrinsic and extrinsic roles of the inflammatory mediators and their receptors (also supported by our single cell sequencing analysis) additional work is needed to understand if and how they contribute to tumor progression in patients with ER mutant tumors. This should include an analysis of MDSC in this setting since they have been described to play an important role in S100A8/A9 function^76,79^. This is also supported by our recent studies showing an enrichment of immune-suppressive macrophages in ER mutant tumors, along with increased expression of interferon regulated genes^80^. Together, these data imply opportunities for immune therapies for patients with ER mutant tumors that should be analyzed further.

We and others^26,27,38^ previously identified genes that have altered expression in *ESR1* mutant cells but are not E2 regulated in WT cells. Here, all six BCK belong to this group of novel, gain-of-function target genes. BCK are not regulated as a result of ligand-mimicking nor *de novo* transactivation by mutant ER, and their expression is strongly and negatively correlated with ER levels. A similar correlation was also observed with P4-induced CK5+ luminal breast cancer cells displaying low ER and PR levels^59^. One possible explanation is that ER, regardless of its liganded status or genotype, serves as a direct epigenetic suppressor that represses BCK expression to maintain luminal identity. For example, it has been shown that ER silences basal, EMT and stem cell related genes by recruiting pivotal methyl-transferases like EZH2 and DNMTs to reshape the DNA and histone methylation landscape^81^. More studies are required to further elucidate the regulatory network between ER and BCKs. Given bi-directional interactions between tumor and stromal cells in BCK regulation, it will be important to perform future studies in improved model systems such as those recently described for analysis of complex regulation of CK14 expression and function^82^.

Assessment of BCK expression revealed that a 50-fold increase in mRNA was reflected in only ~1% cells being positive for BCK protein. This finding is consistent with a previous study showing that P4 stimulation of breast cancer cells caused a 100-fold induction of CK5 promoter activation ultimately translating to 1-10% of cells positive for CK5 protein^59^. In addition, discordance between mRNA and protein of CK7 and CK14 in breast cancer tissue has been documented^83^. It is possible that BCK protein translation in luminal cells is aberrant, resulting in poorly localized or transported protein, consistent with our detection of BCK protein foci rather than the broad distribution pattern over full cytoskeleton similar to what has been previously reported for example for formation of CK17 foci. The discordance in mRNA and protein expression may be due to the cell heterogeneity, with individual cells having high mRNA and protein compared to the negative population, potentially due to heterogenous expression of miRNAs regulating BCK expression^84^. These BCKs positive cells might be pre-selected by multiple genetic and epigenetic cues including but not limited to low ER expression and chromatin loop formation as identified in our study. The discordance between mRNA and protein expression may also help to explain differences in prognosis using mRNA expression profiling like in our study vs IHC in previous studies^85,86^.

We provide evidence to support BCK as emerging biomarkers of *ESR1* mutant breast cancer and its prognosis, yet their direct functional impact remains ambiguous. CK14 positive cells typically lead collective invasion across major subtypes of breast cancer cells^87^, and this is in line with previously identified enhanced cell migration in *ESR1* mutant cells^88^. In addition, as previously described, CK5 positive luminal cells acquire stem-like properties and chemotherapy resistance^47,59^. Importantly, we found several other consistently increased basal marker genes such as interferon-alpha inducible protein 27 (IFI27). Previous studies have reported a role of IFI27 in regulating innate immunity in breast cancer^89^ and cisplatin resistance in gastric cancer^36^. Thus, the “basal-ness” shift might confer several broad functional alterations to *ESR1* mutant tumors.

We identified a PR-orchestrated TAD at the *KRT14/16/17* genomic locus in *ESR1* mutant cells, and we propose that the simultaneous generation of a *de novo* CTCF loop and ER ligand-independent PR overexpression is necessary for *KRT14/16/17* in *ESR1* mutant cells. Intriguingly, knockdown of CTCF selectively increased *KRT14/16/17* mRNA levels whereas knockdown of PR blocked their induction in *ESR1* mutant cells. This unexpected discrepancy may highlight that CTCF binding may simultaneously serve as a transcriptional insulator to restrict *KRT14*/16/17 in an inactive compartment^53,90^. Importantly, data indicates that CTCF knockdown alone is not sufficient to eliminate TAD but instead promotes the formation of new chromatin interactions that alter gene expression^91^. We also unexpectedly found that both PR agonist P4 and PR antagonist RU486 elevated BCK expression, which was inconsistent with previous reported findings where P4 and RU486 exhibited opposite effects in regulating CK5^59^. Given RU486 is well-characterized for its partial agonism, it is possible that *ESR1* mutant cells uniquely express a particular strong PR coactivator that confers the partial agonism of RU486 in this context. Another possibility is that RU486 alternatively stimulates other nuclear receptors such as glucocorticoid^85,92^ or potentially even androgen receptor^93^ to reprogram BCKs expression. The reversed PR pharmacological response in *ESR1* mutant cells is intriguing and warrants future investigation.

Our study discovered a unique aspect of *ESR1* mutant cells and addressed the underlying mechanisms as well as its clinical relevance, albeit with some remaining limitations, such as limited numbers of clinical samples due to inherent difficulties of obtaining metastatic tissues. The enhanced immune infiltration requires additional validation by TIL counting on *ESR1* mutant tumor sections. Confirmation and studies in *in vivo* models should be included into future studies. Our preliminary analysis in a *ESR1* Y541S (mouse ortholog of Y537S mutation) knockin mouse model showed overexpression of BCK at RNA and protein level in mammary tumors^94^. And finally, the *in silico* prediction of enhanced CTCF-driven chromatin loop at the basal cytokeratin gene locus requires confirmation by orthogonal approaches, such as chromosome conformation capture. Nonetheless, our study serves as a robust pre-clinical report uncovering mechanistic insights into *ESR1* mutations and their roles in conferring basal-like feature to ER+ breast cancer and implicates the immune therapeutic vulnerabilities to this subset of patients.

## Materials and methods

Additional details are provided in the Supplementary Materials and Methods section

### Human tissue and blood studies

51 paired primary matched metastatic samples were from DFCI (n=15) and our Women’ s Cancer Research Center (WCRC) (n=36) cohorts as previously reported^95,96^. For all WCRC metastatic samples, *ESR1* mutations status were called from RNA-sequencing. For bone/brain/GI metastatic lesions, *ESR1* mutations status were additionally examined using droplet digital PCR for Y537S/C/N and D538G mutations in *ESR1* LBD region as previously reported^97^. For DFCI cohort, *ESR1* mutations were all called from matched whole exome sequencing^98^.

For the study of patients’ blood, all patients provided written informed consent and all procedures were approved by the University of Pittsburgh Institutional Review Broad (PRO17080172). 18 patients diagnosed with late-stage metastatic ER+ breast cancer were recruited. Procedure to identify hotspot *ESR1* mutations has been previously described by us^99^.

### Cell culture

Establishments of rAAV-edited (Park lab)^27^, CRISPR-Cas9-edited (Gertz^38^ and Ali^25^ lab) and CRISPR-Cas9-edited T47D cells^27^ were reported previously. ZR75-1 (CRL-1500), MDA-MB-134-VI (HTB-23), MDA-MB-330 (HTB-127) and MDA-MB-468 (HTB-132) were obtained from the ATCC. Development of BCK4 cells has been previously reported^100^.

### S100A8/S100A9 heterodimer ELISA

Human S100A8/S100A9 heterodimer amounts in human plasma samples were quantified using S100A8/S100A9 heterodimer Quantikine ELISA kit (R&D System, DS8900) following the manufacture protocol. All plasma samples were first diluted in calibration buffer with 1:50 ratio and loaded into antibody-coated plate.

### Chromatin-immunoprecipitation (ChIP) and sequencing analysis

ChIP was performed as previously described ^51^. ChIP-seq reads were aligned to hg38 genome assembly using Bowtie 2.0 ^101^, and peaks were called using MACS2.0 with p value below 10E-5 ^102^. We used DiffBind package ^103^ to perform principle component analysis, identify differentially expressed binding sites and analyze intersection ratios with other data sets. Heatmaps and intensity plots for binding peaks were visualized by EaSeq. Annotation of genes at peak proximity was conducted using ChIPseeker ^104^, taking the promoter region as +/− 3000 bp of the transcriptional start site (TSS) and 50kb as peak flank distance.

### RNA sequencing analysis

RNA sequencing data sets were analyzed using R version 3.6.1. Log2 (TPM+1) values were used for the RNA-seq of Oesterreich *ESR1* mutant cell models and TMM normalized Log2(CPM+1) values were used for Gertz RNA-seq data. TCGA reads were reprocessed using Salmon v0.14.1^105^ and Log2 (TPM+1) values were used. For the METABRIC data set, normalized probe intensity values were obtained from Synapse. For genes with multiple probes, probes with the highest inter-quartile range (IQR) were selected to represent the gene. For pan-breast cancer cell line transcriptomic clustering, 97 breast cancer cell line RNA-seq data were reprocessed using Salmon and merged from three studies^34–36^, batch effects were removed using “removeBatchEffect” function of “limma^106^” package. Gene set variation analysis was performed using “GSVA” package^107^. Survival comparisons were processed using “survival” and “survminer” packages^108^ using Cox Proportional-Hazards model and log-rank test. Data visualizations were performed using “ggpubr^109^”, “VennDiagram^110^” and “plot3D^111^”.

For the single cell RNA seq analysis, two fresh bilateral bone metastases (BoMs) were collected from a patient initially diagnosed with ER+ primary breast cancer, dissociated into single cells and a cell suspension with at least 70% viability was submitted for library preparation using 10X genomics chromium platform (V3.0 chemistry) (Ding et al, manuscript in preparation). 6,000 cells were targeted for each BoM, and the final libraries were sequenced at a depth of 67,000 reads per cell using NOVAseq.

### Tumor Mutation Burden Analysis

Tumor mutation burden (TMB) calculation was performed as previous described^112^. Briefly, TCGA mutation annotation files from 982 patients were downloaded from FireBrowse and mutation subtypes were summarized using “maftool” package^113^. Mutations subtypes were classified into truncated (nonsense, frame-shift deletion, frame-shift insertion, splice-site) and non-truncated mutations (missense, in-frame deletion, in-frame insertion, nonstop). TMB was calculated as 2X Truncating mutation numbers + non-truncating mutation numbers.

### Generation of Gene Sets

For Sorlie et al., the original set of intrinsic genes were downloaded from Stanford Genomics Breast Cancer Consortium (http://genome-www.stanford.edu/breast_cancer/). 453 genes were annotated from 553 probes. Expression of these 453 genes were examined in 33 luminal and 39 basal breast cancer cell lines. Significantly higher (FDR<0.01) intrinsic genes in basal or luminal cells were called as basal (n=75) or luminal (n=68) markers in Sorlie gene sets. For the TCGA gene set, differentially expressed genes were called between basal and luminal A or basal and luminal B ER+ tumors using raw counts. The top 200 increased genes of these two comparisons were further intersected. Overlapped DE genes in basal (n=164) and luminal (n=139) tumors were called as TCGA gene sets. For CTCF gene signature establishment, a previous RNA-seq data set on MCF7 cells with or without CTCF knockdown was downloaded and analyzed^52^, top 100 downregulated genes with CTCF knockdown were used as the CTCF gene signature.

### Chromatin interaction data analysis

CTCF ChIA-PET data were downloaded from GSE72816. Chromatin linkages were visualized on 3D genome browser (http://promoter.bx.psu.edu/hi-c/) after processed with ChIA-PET tool^114^. Confident TAD boundaries were defined by the colocalization of CTCF and cohesion complex subunits together with called chromatin interactions.

## Data Availability

ER ChIP-seq data from MCF7 *ESR1* mutant cell model was deposited in Gene Expression Omnibus with accession number of GSE125117. MSigDB curated gene sets were downloaded from GSEA website (http://software.broadinstitute.org/gsea/msigdb/index.jsp). RNA-seq data and clinical information from TCGA and METABRIC were obtained from the GSE62944 and Synapse software platform (syn1688369) respectively. TCGA biospecimen immune profile data were downloaded from Saltz et al^65^. TCGA mutation annotation format (MAF) files and methylation data were downloaded from FireBrowse website (http://firebrowse.org/). Complete RNA-Seq data for the DFCI metastases samples will be published separately. RNA-Seq data from the WCRC cohorts are available at Lee-Oesterreich Lab Github repository (https://github.com/leeoesterreich). All the raw data and scripts are available upon request from the corresponding author. Sources of all public available data sets used in this study are summarized in Supplementary Table S10.

## Acknowledgements

We thank Dr. Peilu Wang for her contribution to earlier *ESR1* mutant-studies in the Lee-Oesterreich group. We also thank Corrine Farrell and Jian Chen for some technical support of the project. This project used the University of Pittsburgh Pitt Biospecimen Core (PBC), supported in part by award NIH grant P30CA047904. The authors would like to thank the patients who contributed samples to the tissue bank as well as all the clinicians and staff for their efforts in collecting tissues.

## Author Contributions

Z.L., J.M.A., A.V.L. and S.O. conceived and designed the study. Z.L., Y.W., A.B. and K.D. designed, performed and analyzed experiments. Z.L., A.B., N.M.P. and K.D. performed bioinformatic analysis. J.M.A., L.M. and M.R. contributed to clinical sample collection and intellectual input. N.W. provided extended RNA-seq data set (DFCI) from clinical specimens and intellectual input. Z.L., A.V.L., S.O., C.A.S., J.K.R., W.J.M. and J.G. contributed to data interpretation and provided additional intellectual input. L.B., S.A. and J.G. provided additional cell models for this study and intellectual input. Y.F., L.Z. and G.C.T. provided and validated biostatistical approaches of all the analysis. Z.L., A.V.L. and S.O. developed the figures and the manuscript. All the authors reviewed and agreed with the contents of the manuscript.

