## Supplementary Figures for "*ESR1* mutant breast cancers show elevated basal cytokeratins and immune activation"

Supplementary Figure S1

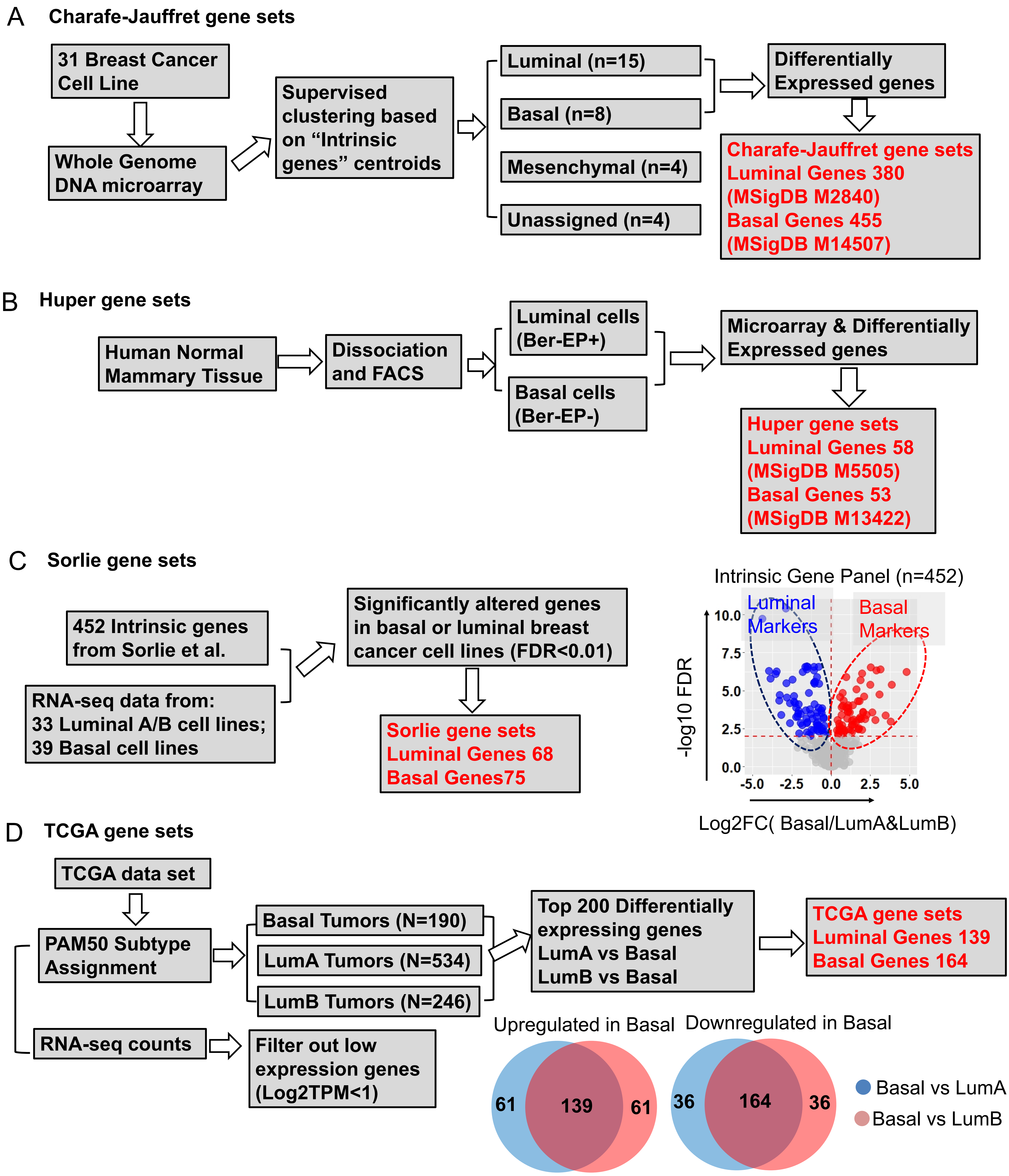


**Supplementary Figure S1. Schematic flow charts of the generation process of the four pairs of luminal/basal gene sets. (Related to Fig. 1)**

A) Charafe-Jauffret gene set was obtained based on a microarray data of 31 breast cancer cell lines after supervised clustering and subtype assignment. Differentially expressed genes from 15 luminal and 8 basal breast cancer cells derived 380 and 455 luminal and basal genes.

B) Huper gene set was computated based on microarray data of flow cytometry sorted basal and luminal epithelial cells from a normal human mammary tissue. This gene set contains 58 luminal and 53 basal genes.

C) Sorlie gene set was derived based on the differentially expressed genes between 39 basal and 33 luminal breast cancer cell lines among the 452 intrinsic gene panel reported by Sorlie et al. The volcano plot on the right panel represents the differentially gene expression distribution of the 452 genes, genes with FDR<0.01 were highlighted and used for the gene sets.

D) For the TCGA gene set, differentially expressed genes(FDR<0.01) were called between basal and luminal A or basal and luminal B ER+ tumors using raw counts after filtering out low expression genes (maximum Log2 TPM expression across all samples <1). The top 200 increased genes from these two comparisons were further intersected (Venn diagram). Shared upregulated (n=139) or downregulated (n=164) DE genes between basal-to-LumA and basal-to-LumB comparisons tumors were called as TCGA gene sets.

### Supplementary Figure S2

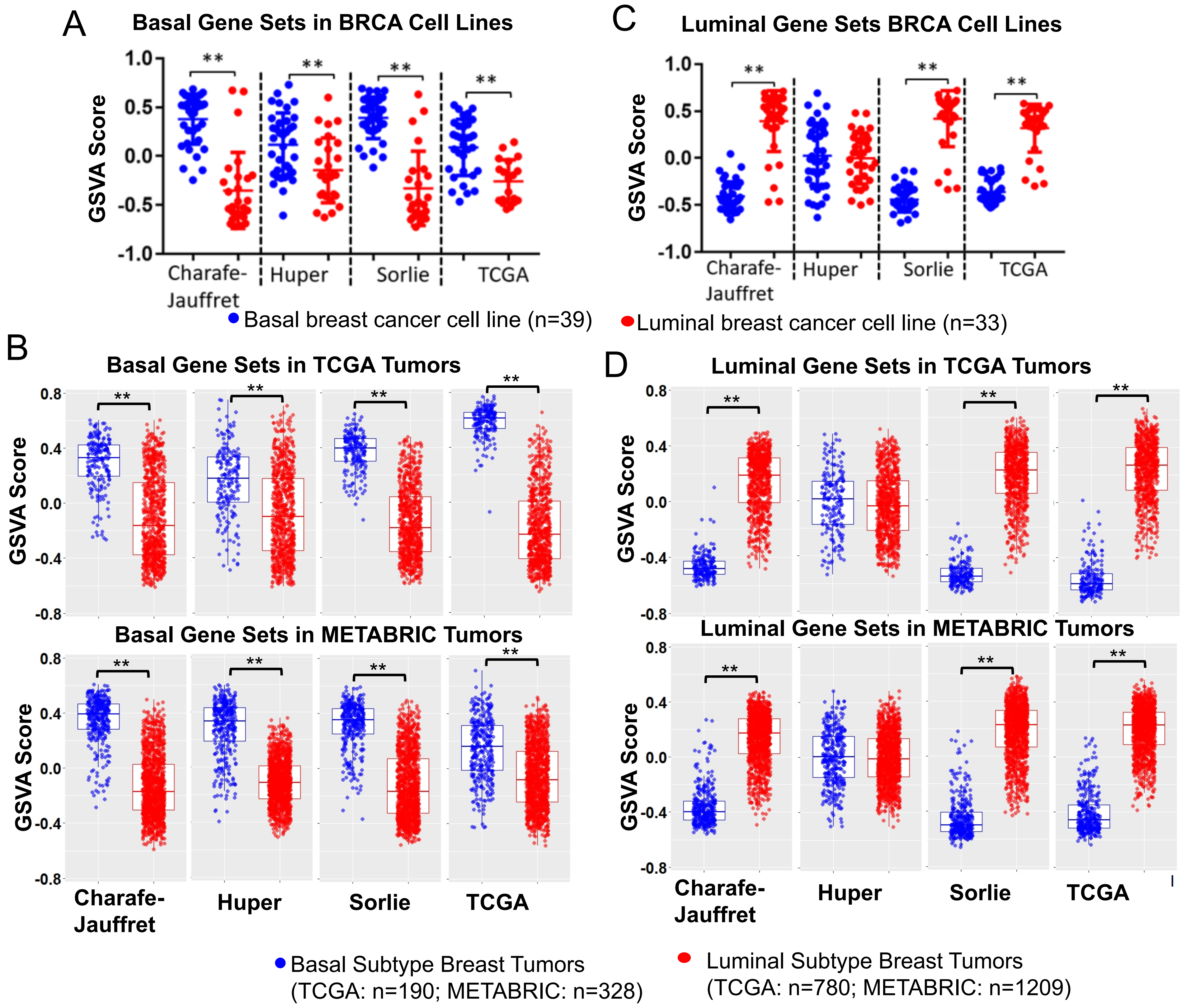


**Supplementary Figure S2. Quality control of the four pairs of luminal/basal gene sets in breast cancer cell lines and tumors. (Related to Fig. 1)**

A) and C) Dot plots showing GSVA score of the four pairs of basal (A) /luminal (C) gene sets enrichment in luminal (n=33) and basal (n=39) breast cancer cells as quality controls. Each plot represents mean $\pm$ SD from GSVA score. Mann Whitney U test was used within each group (** $p<0.01$).

B) and D) Box plots showing GSVA score of the four pairs of basal (B) /luminal (D) gene sets enrichment in luminal and basal breast cancer tumors from TCGA (n=190 basal, n=780 luminal) and METABRIC (n=328 basal, n=1209 luminal) as quality controls. Each plot represents median $\pm$ SD from GSVA score. Mann Whitney U test was used within each group. ( ** $p<0.01$).

### Supplementary Figure S3

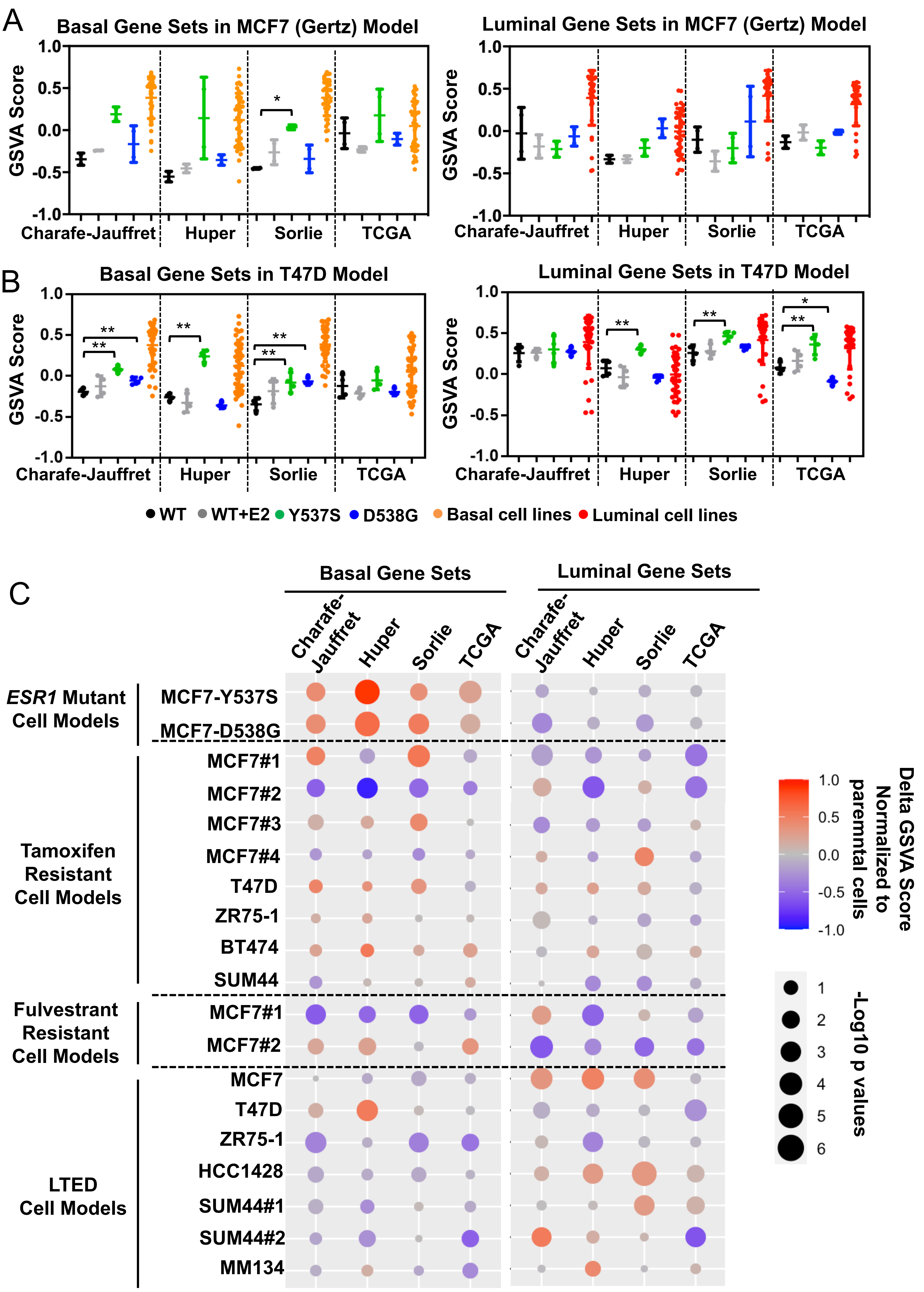


**Supplementary Figure S3. *ESR1* mutant cell models but not other endocrine resistant models show increased basal marker enrichment. (Related to Fig. 1)**
A) and B) Dot plots showing GSVA score of the four pairs of basal/luminal marker gene sets enrichment in MCF7 (Gertz) (A) and T47D (B) genome-edited *ESR1* mutant cell models. Two individual clones were used as biological replicates for Gertz model, while four biological replicates were used for T47D cell model. Enrichment scores in luminal (n=33) and basal (n=39) breast cancer cell lines were used as positive controls. Dunnett's test was used within each group. (* p<0.05, ** p<0.01)

C) Graphic view of delta GSVA enrichment score of 19 different endocrine resistant breast cancer models normalized to their corresponding parental cell counterparts. Color scale represents changes of enrichment scores and dot size shows significance. Data sets were downloaded and analyzed from GEO. (Tamoxifen resistant models: GSE128458, GSE104985, GSE111151, GSE26459 and GSE106681; Fulvestrant resistant models: GSE118713 and GSE104985; LTED models: GSE75971 and GSE116744). Deatailed sources information can be found in Supplementary Table S10.

Supplementary Figure S4

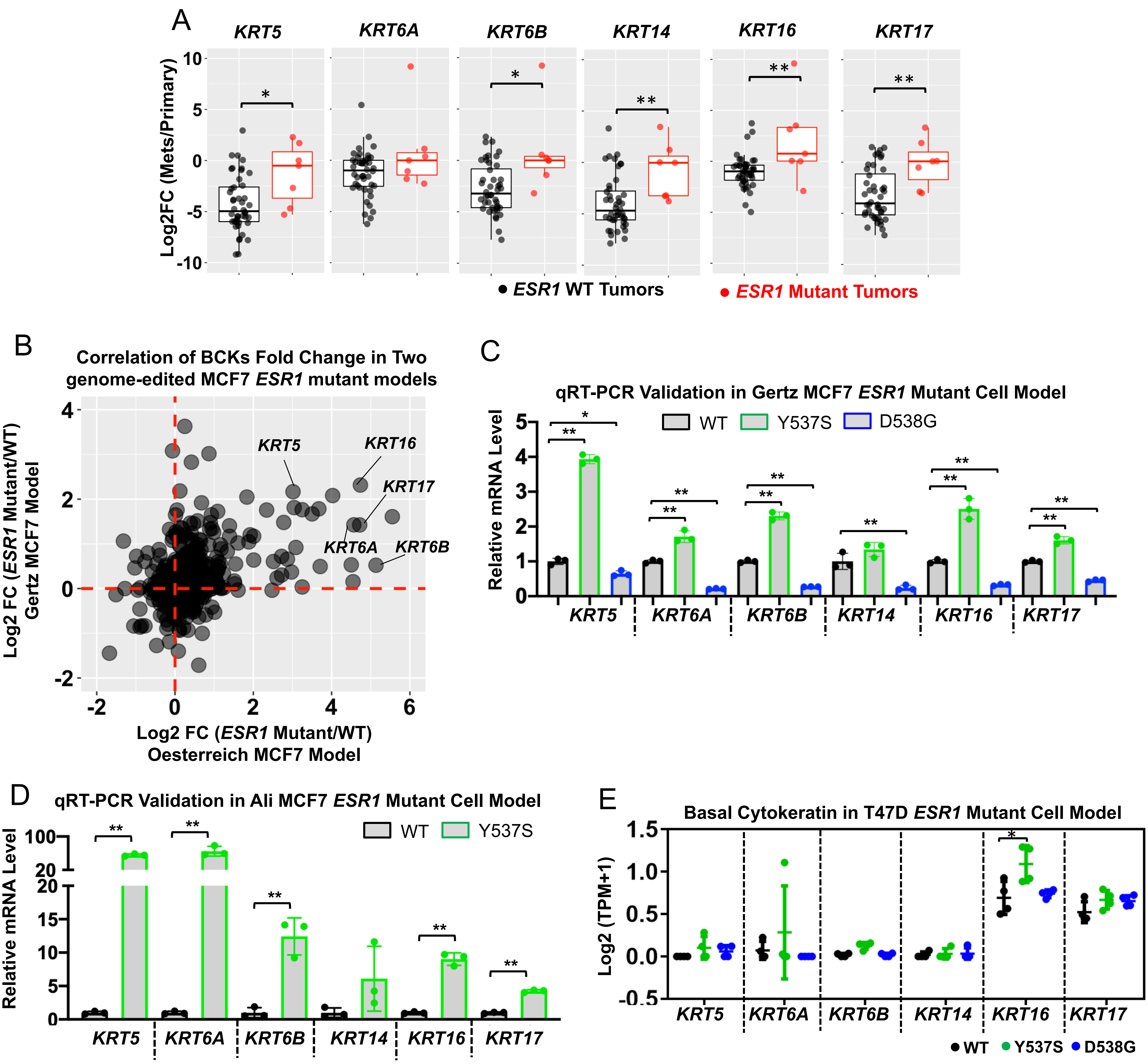


**Supplementary Figure S4. Basal cytokeratins are increased in other genome-edited *ESR1* mutant cell models. (Related to Fig. 2)**

A) Box plots representing the six basal cytokeratin expression in primary-matched paired metastatic tumor samples. Log2 (CPM+1) values were used for calculation. Expressional fold changes in each metastatic tumor were normalized to the matched primary tumor. Mann-Whitney U test was performed to compare the expression between *ESR1* WT (N=44) or *ESR1* mutant (N=7) paired tumors. (* $p<0.05$; ** $p<0.01$)

B) Two dimensional plot showing the correlation of basal marker gene fold changes in Oesterreich and Gertz MCF7 genome-edited Y537S/D538G cells (normalized to WT vehicle group). Increased basal cytokeratins were labelled with gene names. (* $p<0.05$; ** $p<0.01$)

C) and D) Bar graphs representing qRT-PCR measurement of *KRT5/6A/6B/14/16/17* mRNA levels in the Gertz (C) and Ali (D) MCF7 WT and *ESR1* mutant cells. ΔΔCt method was used to analyze relative mRNA fold changes normalized to WT cells and *RPLP0* levels were measured as the internal control. Each bar represents mean ± SD with three biological replicates. These experiments were done once for each. Dunnett's test (C) and student's t test (D) were used to compare BCKs expression levels between WT and mutants respectively. (** $p<0.01$)

E) Dot plot representing all six basal cytokeratins expression in the Oesterreich T47D *ESR1* mutant cell models.  Each dots represent the Log2 (TPM+1) value from RNA-seq with four biological replicates. Dunnett's test was applied. (* $p<0.05$)

Supplementary Figure S5

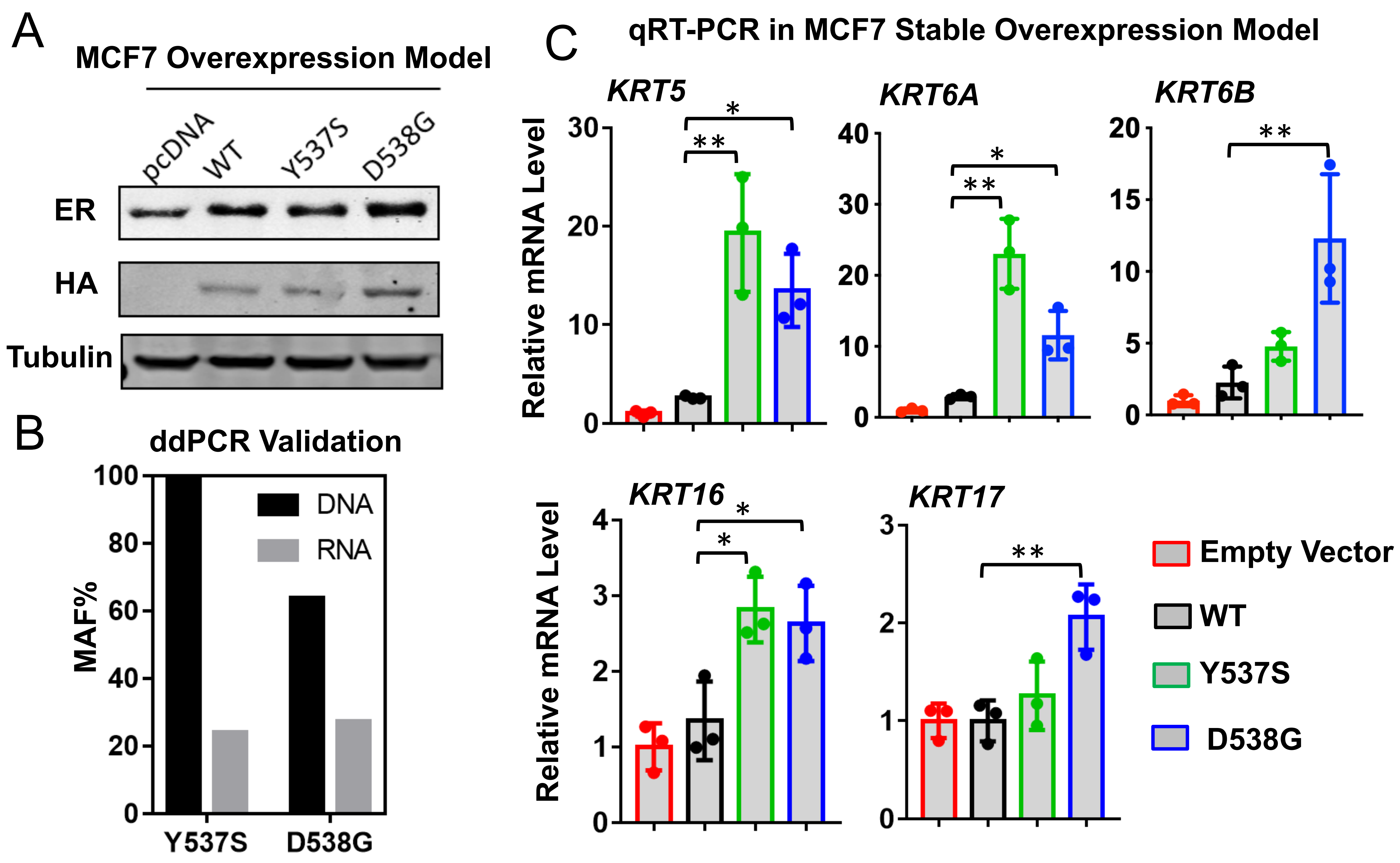


**Supplementary Figure S5. Basal cytokeratins are increased in stablely overexpressed *ESR1* mutant cell models. (Related to Fig.2)**

A) Immunoblot validation of MCF7 *ESR1* mutant overexpression cells. Total ER and overexpressed ER (HA-tagged) were detected and tubulin was used as loading control. This experiment was done once.

B) ddPCR validation of mutation allele frequencies in Y537S and D538G overexpression cell model at both DNA (gDNA) and RNA (cDNA) levels.

C) Bar graphs representing qRT-PCR measurement of *KRT5/6A/6B/16/17* mRNA levels in MCF7 overexpression *ESR1* mutant cell models. ΔΔCt method was used to analyze relative mRNA fold changes normalized to empty vector cells and *RPLP0* levels were measured as the internal control. Each bar represents mean ± SD with three biological replicates. Data were from one representative experiment of two independent repeats. Dunnett's test was used to compare the gene expression of each *ESR1* mutant group to WT cells. (* p<0.05, ** p<0.01).

Supplementary Figure S6

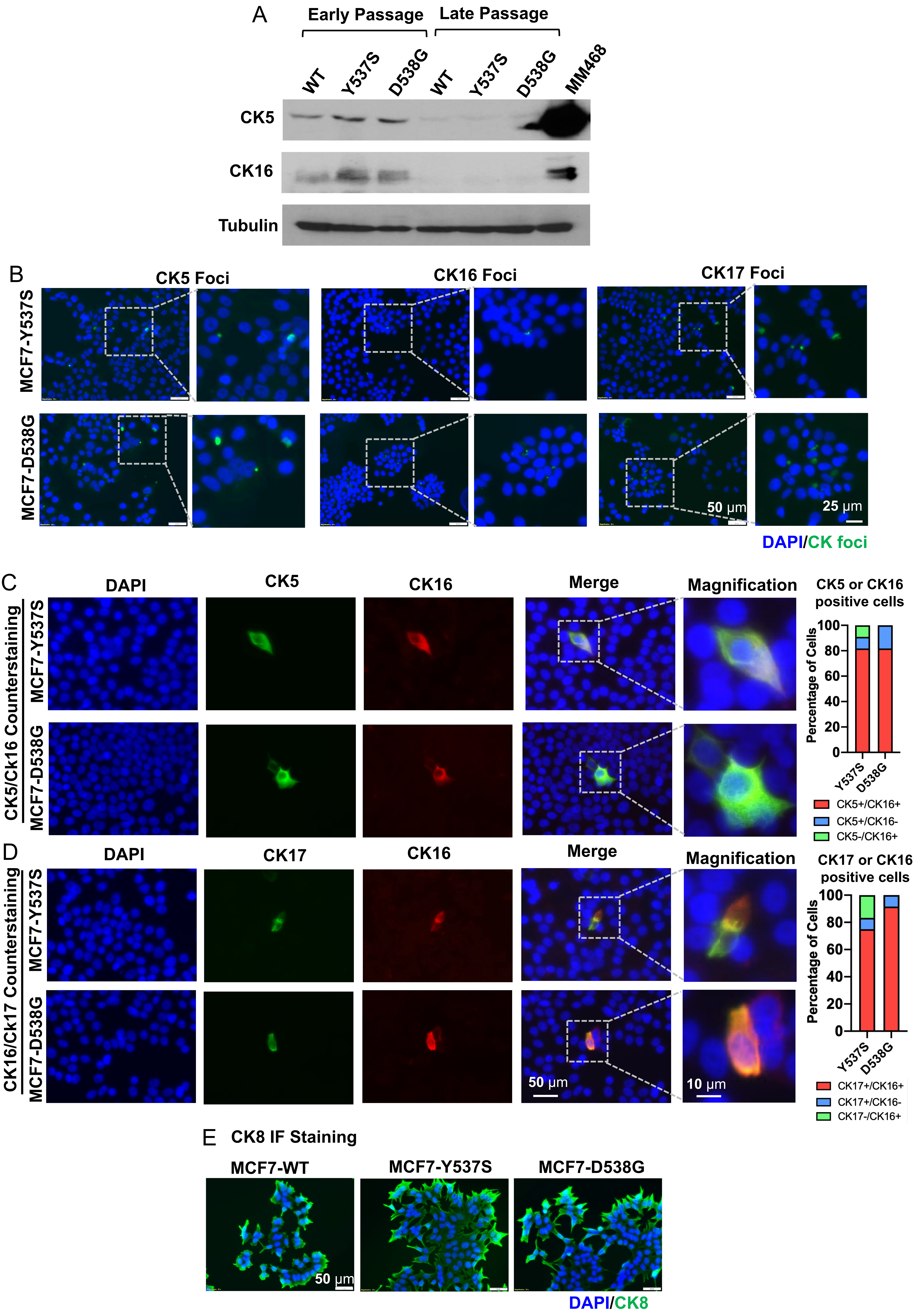

**Supplementary Figure S6. Basal cytokeatins are heterogeneously expressed in *ESR1* mutant cells. (Related to Figure 2)**

A) Immunoblot validation of CK5 and CK16 expression MCF7 *ESR1* mutant cells in low (P6-P8) and high (P30-P32) passages. Tubulin was used as loading control. This experiment was done once.

B) Representative images of CK5/16/17 foci in MCF7 *ESR1* mutant cells. Images were taken under 20x magnification. Specific regions with foci detected were further zoomed in as magnificated views. Data were from one representative of three independent replicates.

C) and D) Left panel: Representative images of CK5/CK16 (C) and CK5/CK17 (D) counterstaining in MCF7 *ESR1* mutant cells. Images were taken under 20x magnification. Specific regions with CK+ cells were further zoomed in as magnificated views. Right panel: Stacked plots representing the quantification of percentage of cells with double CK subtype positivity (red) or single CK subtype positivity (blue/green) among all CK+ cells (n=11 for CK5/CK16 costaining quantficiation, n=12 for CK16/CK17 costaining quantification). This experiment was done once.

E) Representative images of luminal cytokeratin CK8 staining in MCF7 WT and *ESR1* mutant cells. Images were taken under 20x magnification. This experiment was done once.

Supplementary Figure S7

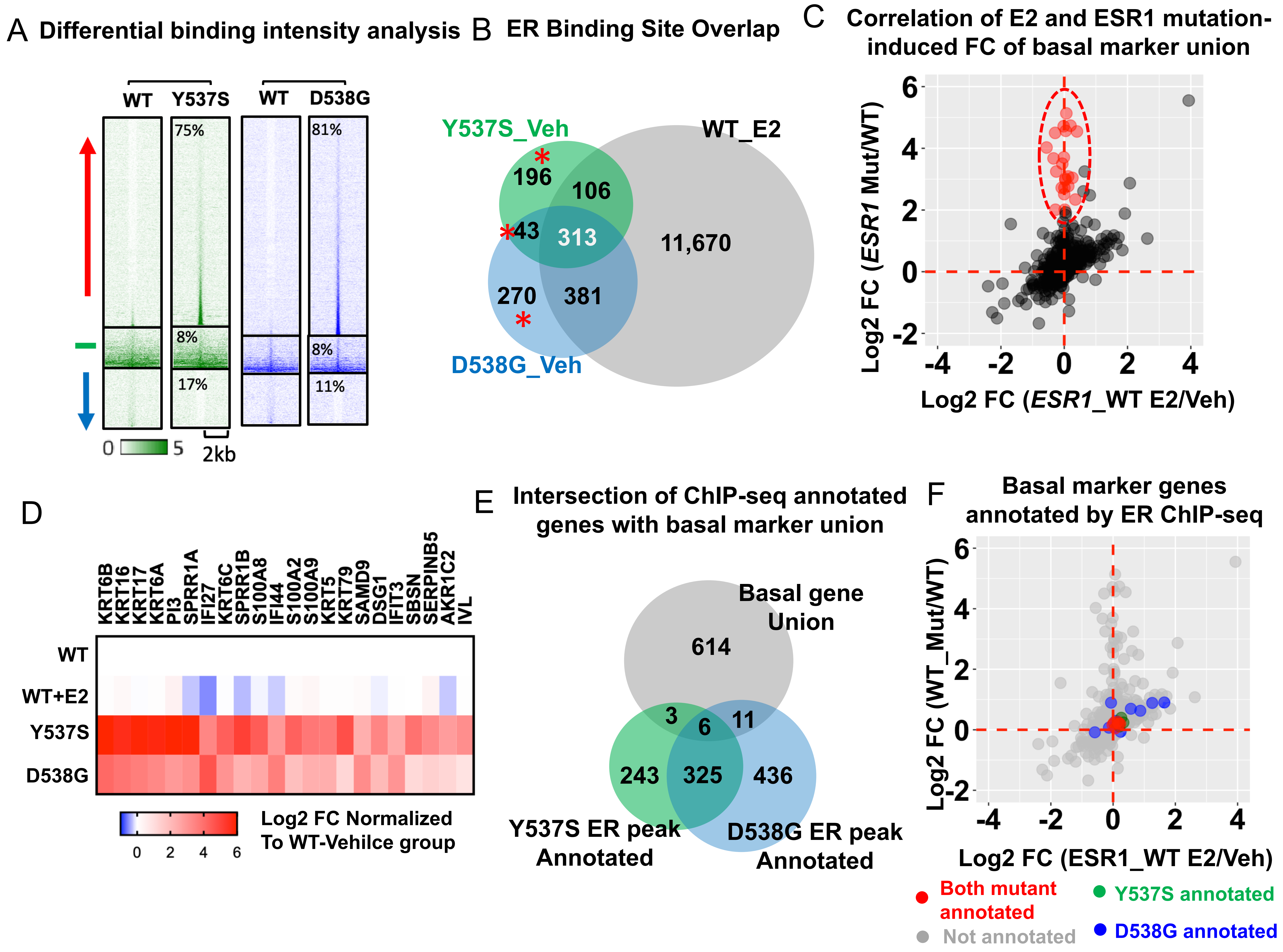


**Supplementary Figure S7. ChIP-seq profiling of mutant ER cistrome reveals limited regulation on basal marker gene from ER genomic binding. (Related to Fig.3)**

A) Heatmaps of differential ER binding intensities in Y537S, D538G mutants compared to WT ER in a pairwise manner, shown in a horizontal window of ± 2kb from the peak center. The pairwise comparison between WT and mutant samples were performed to calculate the fold change (FC) of intensities and the binding sites were sub-classified into sites with increased intensity (FC>2, red arrow), decreased intensity (FC<-2, blue arrow), and non-changed intensity (-0.2<FC<0.2, green line). Percentage of each subgroups were labelled on the heatmaps respectively.

B) Venn Diagrams showing the occupancy intersection between WT-E2, Y537S-vehicle and D538G-vehicle groups in MCF7 cell model. *De novo* mutant ER peaks were labelled with asterisk symbols.

C) Two dimensional plot showing the correlation of basal gene fold changes of MCF7 *ESR1* mutant (average FC of Y537S and D538G) cells and E2 stimulation in WT cells. 22 skewed basal genes were highlighted in red (Log2FC>2 in mutants and <0.5 in WT-E2 group).

D) Heatmap representing the fold changes of the 22 unqiue *ESR1* mutant-regulated basal genes (highlighted in Figure 7C) in WT-E2 and two mutant groups normalized to WT-vehicle group using RNA-seq data (GSE89888).

E) Venn digram showing interaction of Y537S (n=577) and D538G (n=778) ER peak annotated genes to the union of basal genes (n=634) collection. Annotated genes were called within -/+ 50 kb range of each ER peak.

F) Two dimensional plot showing the correlation of basal marker gene fold changes of MCF7 *ESR1* mutant (average FC of Y537S and D538G) cells and E2 stimulation in WT cells. Genes that were annotated by ER ChIP-seq data in Y537S (green), D538G (blue) and both mutants (red) were labelled accordingly.

Supplementary Figure S8

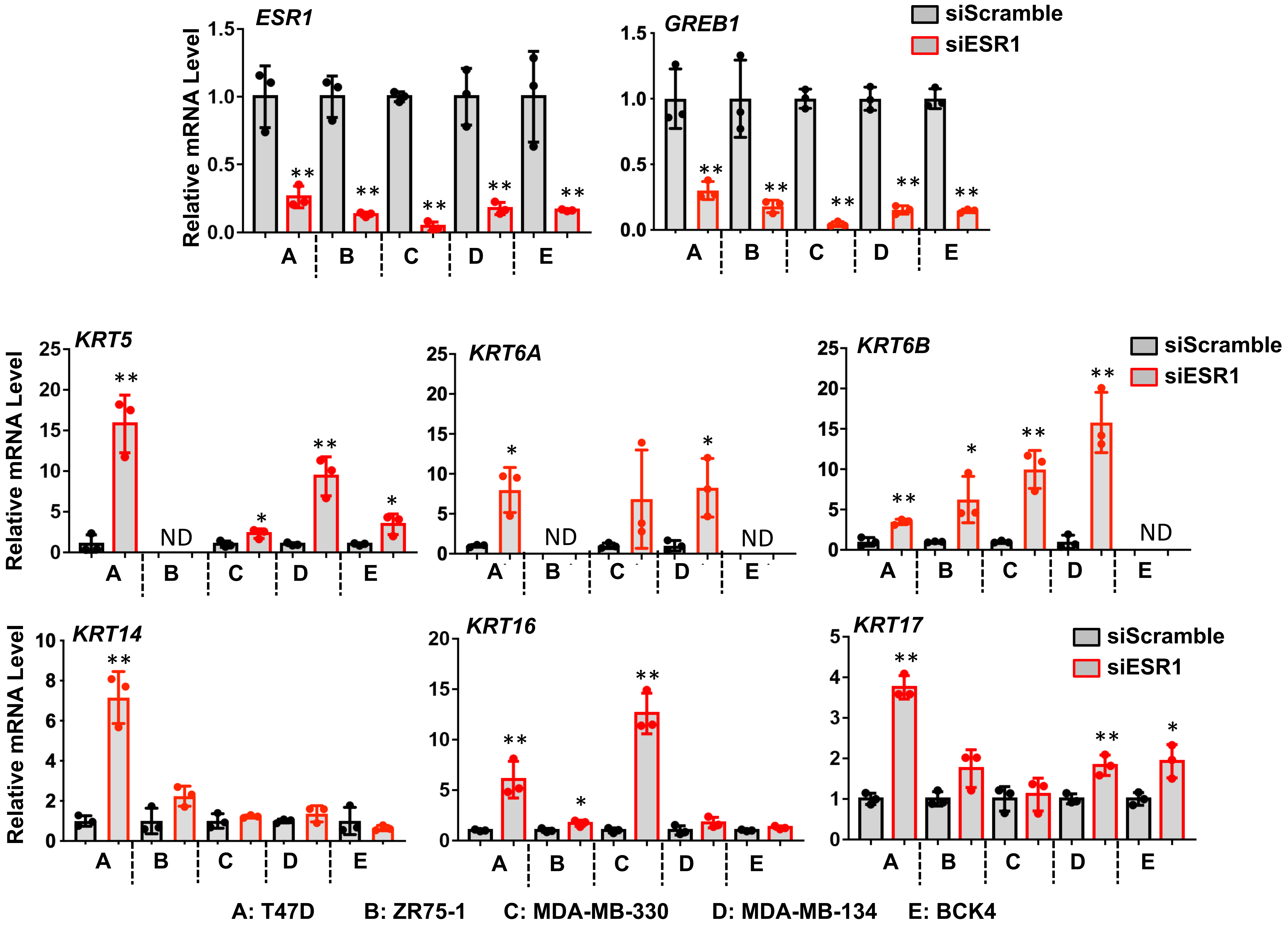


**Supplementary Figure S8. Knockdown of *ESR1* generally increased basal cytrokeratin expression in ER+ breast cancer cell lines. (Related to Fig. 3)**

Bar graphs representing qRT-PCR measurement of *ESR1*, *GREB1* and *KRT5/6A/6B/14/16/17* mRNA levels in five ER+ breast cancer cells with siRNA knockdown of *ESR1* for 7 days. ΔΔCt method was used to analyze relative mRNA fold changes normalized to each siScramble groups and *RPLP0* levels were measured as the internal control. Each bar represents mean ± SD with three biological replicates. Student's test was used to compare the gene expression between scramble and knockdown groups. This experiment was done once. Genes with undetectable values were indicated as ND. (* $p<0.05$, ** $p<0.01$).

Supplementary Figure S9

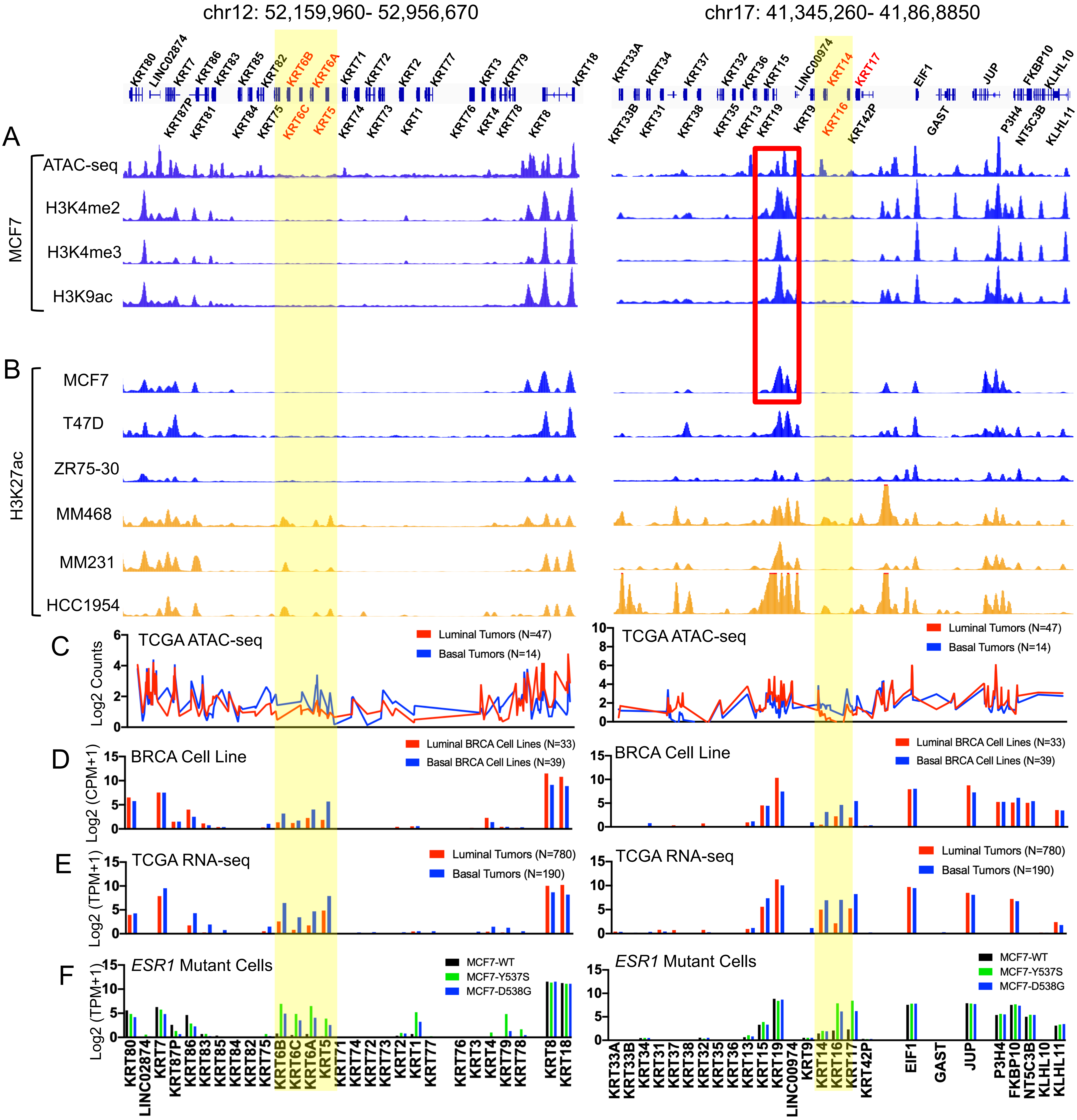


**Supplementary Figure S9. Epigenetic landscapes of *KRT5/6A/6B* and *KRT14/16/17* loci in basal and luminal cells and tumors. (Related to Fig. 4)**

Genomic track illustrations of *KRT5/6A/6B* and *KRT14/16/17* proximal regions with active histone modifications ChIP-seq (A) in MCF7 and H3K27 acetylation ChIP-seq in 3 luminal and 3 basal breast cancer cell lines (B). Data set were visualized at WashU Genome Browser based on public available data sets from GEO (GSE102441, GSE69112, GSE65201 and GSE29069) and ENCODE (ENCSR875KOJ, ENCSR958MIB, ENCSR056UBA and ENCSR752UOD). Y axis represent the signal intensity of ChIP-seq data sets. TCGA ATAC-seq (C) and RNA-seq log2 (TPM+1) values (E) were compared between luminal and basal tumors with specific numbers labelled in the plots. In addition, proximal gene expression were compared in 33 luminal and 39 basal breast cancer cell lines (D) and MCF7 *ESR1* mutant cell models (F) using Log2 (CPM+1) and Log2 (TPM+1) values respectively based on RNA-seq data. *KRT5/6A/6B* and *KRT14/16/17* loci were highlighted in light yellow. The super enhancer region featured by active histone modification at *KRT14/16/17* loci was highlighted in a red frame.

Supplementary Figure S10

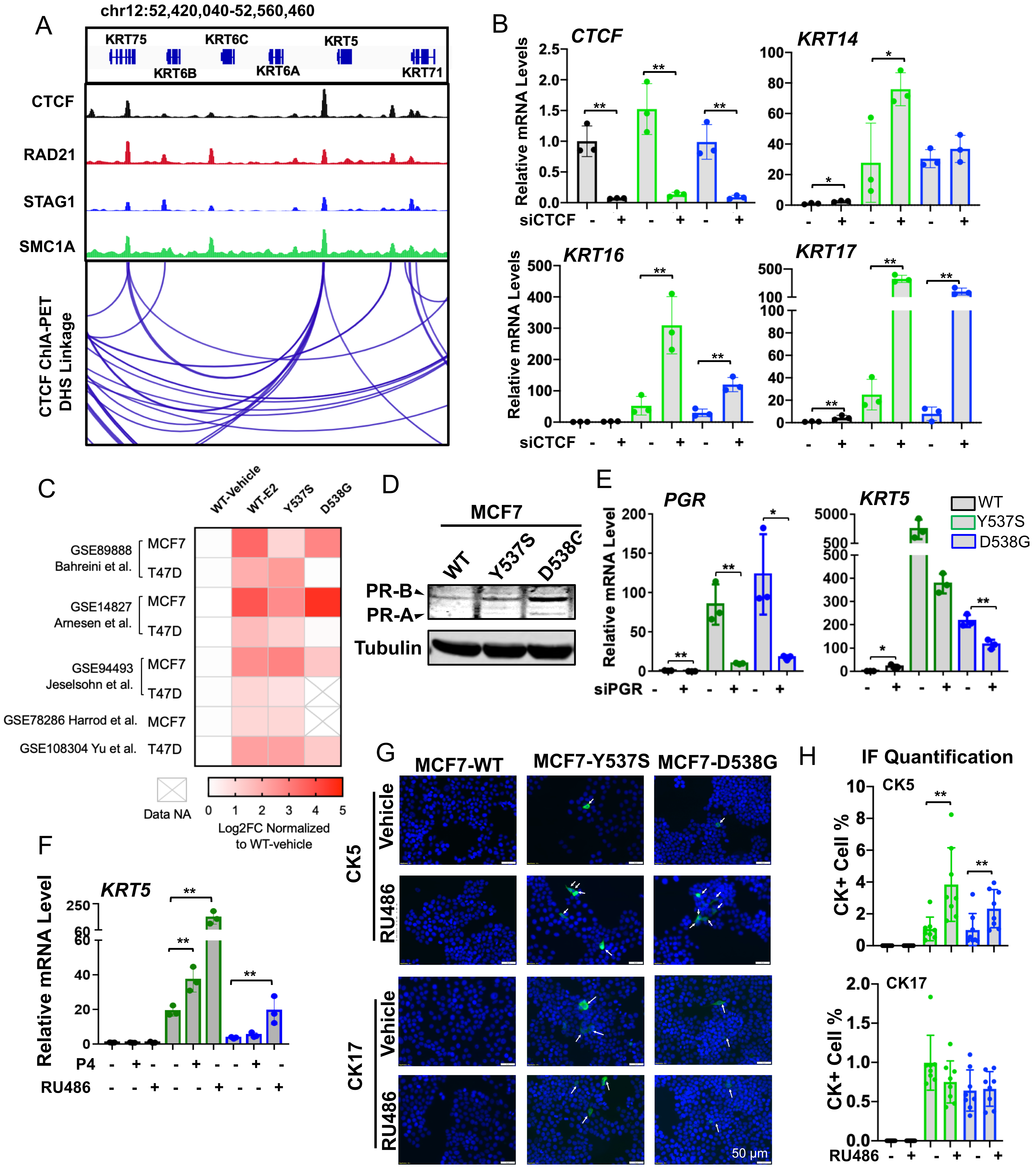

**Supplementary Figure 10. BCKs are associated with a progesterone receptor binding enhancer orchestrated TAD in *ESR1* mutant cells. (Related to Fig.4)**

A) Upper panel: Genomic track illustrating the CTCF/cohesion complex binding at *KRT5/6A/6B* proximal genomic region in MCF7 cells. CTCF and RAD21 ChIP-seq were downloaded from ENCODE (ENCSR560BUE and ENCSR703TNG). STAG1 and SMC1A ChIP-seq data were from GEO (GSE25021 and GSE76893). Y-axis represents signal intensity of each track. Include genomic coordinates. Lower panel: CTCF-driven chromatin loops visualized using a CTCF ChIA-PET data set in MCF7 cells (GSE72816) at the 3D Genome Browser platform. Each linkage represents a chromatin loop.
B) Bar graphs showing qRT-PCR measurement of *CTCF*, *KRT14, 16* and *17* mRNA levels in MCF7 *ESR1* WT and mutant cells with siRNA knockdown of *CTCF* for 7 days. ΔΔCt method was used to analyze relative mRNA fold changes normalized to WT cells (siScramble group) and *RPLP0* levels were measured as the internal control. Each bar represents mean $\pm$ SD with three biological replicates. Student's test was used to compare the gene expression between scramble and knockdown groups. This experiment was done once. (* $p<0.05$, ** $p<0.01$)

C) Heatmap representing *PGR* gene expression fold change in WT+E2 and *ESR1* mutant groups from eight different *ESR1* mutant cell models with public available RNA-seq data sets .

D) Immunoblot detection PR in MCF7 *ESR1* mutant cells. Tubulin was used as a loading control. This experiment was done once.

E) Bar graphs showing qRT-PCR measurement *PGR* and *KRT5* mRNA levels in MCF7 *ESR1* WT and mutant cells with siRNA knockdown of *PGR* for 7 days. Method was the same as described in B. Data is from one representative of three independent replicates. (* $p<0.05$, ** $p<0.01$)

F) Bar graphs showing qRT-PCR measurement *KRT5* mRNA levels in MCF7 *ESR1* WT and mutant cells with 0.1% EtOH, 100 nM P4 and 1 μM RU486 treatment for 3 days. Method was the same as described in B. Data is from one representative of three independent replicates. (* $p<0.05$, ** $p<0.01$)

G) Representative images of immunofluorescence staining on CK5 and CK17 in MCF7 WT and *ESR1* mutant cells under 1 μM RU486 treatment for 3 days. Images were taken under 20x magnification. CK+ cells are pointed with white arrows.

H) Bar plots quantifying the percentages of CK positive cells of each group. Each bar represents mean $\pm$ SD from eight different regions combining from two independent experiments. Student's t test was used to compare BCKs positive cell percentage in the presentce and absence of the treatment. (* $p<0.05$, ** $p<0.01$)

Supplementary Figure S11

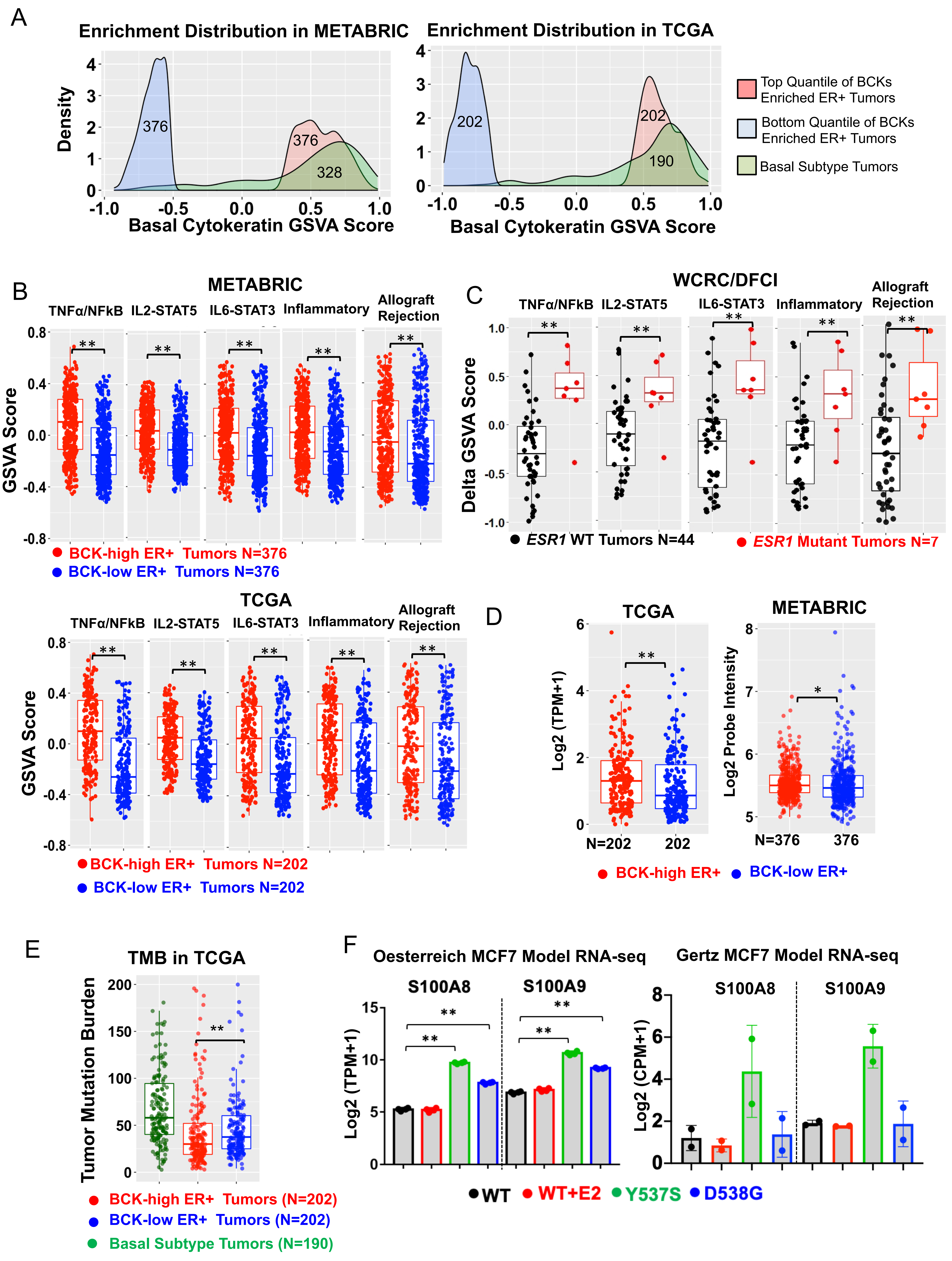

**Supplementary Figure 11. Basal cytokeratins are associated with enhanced immune activation in ER+ tumors. (Related to Fig. 5 and 6)**

A) Density plots showing the expressional distributon of BCKs GSVA scores in TCGA and METABRIC ER+ tumors for the bottom (blue) and top (red) quantile used for downstream analysis. BCKs enrichment in basal breast cancers (green) was set as control. Specific numbers of each subpopulation were indicated in the plots.

B) and C) Box plots showing the enrichment scores of the five intersected immune-related pathways between BCK-high (n=202 TCGA; n=376 METABRIC) and low (n=202 TCGA; n=376 METABRIC) subsets in ER+ tumors in METABRIC (B, Top panels) and TCGA (B, Lower panel), and between intra-patient primary tumor-paired *ESR1* mutant (n=7) and WT (n=44) metastatic lesions (C). Mann Whitney U test was used for each comparison. (**p<0.01)

D) Box blots representing *PDCD1* expression between BCKs-high (n=202 TCGA; n=376 METABRIC) and low (n=202 TCGA; n=376 METABRIC) ER+ tumors in TCGA and METABRIC. Log2 (TPM+1) from TCGA RNA-seq and log2 normalized probe intensity from METABRIC were used for plot. Mann Whitney U test was used for each comparison. (*p<0.05, **p<0.01)

E) Box plot showing comapresion of tumor mutation burdens in TCGA basal subtype tumors (n=190) and BCKs high (n=202) and low (n=202) ER+ tumors. Tumor mutation burdens were calculated as 2X truncating mutation numbers + non-truncating mutation numbers of each tumor. Mann Whitney U test was applied. (**p<0.01)

F) Bar graphic view of S100A8 and S100A9 gene expression based on RNA-seq data of Oesterreich (n=4, Log2 (TPM+1)) and Gertz (n=2, Log2 (CPM+1)) MCF7 *ESR1* mutant cells models. Dunnett’s test was utilized for the Oesterreich model.  (** p<0.01)
