## Supplementary Materials and Methods for "*ESR1* mutant breast cancers show elevated basal cytokeratins and immune activation"

***ESR1* mutation detection**

Hotspot *ESR1* mutation identification procedure was previously described by us^1^. In brief, blood samples were collected in EDTA tubes (BD, #367856) and cfDNA was isolated from plasma samples using Qiagen circulating nucleic acid kit (#55114). *ESR1* ligand binding domain was pre-amplified in cfDNA and the products were subjected to droplet digital PCR detection with Y537S/C/N and D538G probes.

**Cell culture**

Establishments of rAAV-edited (Park lab)^2^ , CRISPR-Cas9-edited (Gertz^3^ and Ali^4^ lab) and CRISPR-Cas9-edited T47D cells^2^ were reported previously. Individual clones were maintained in DMEM, supplemented with 10 % FBS, 100 μg/mL penicillin and 100 mg/mL streptomycin, at 37 °C in a humidified incubator with 5% CO2. Mutation allele frequencies were confirmed using ddPCR. For all experiments, hormone deprivation was performed unless stated otherwise, cells were maintained in phenol-red-free IMEM (Gibco, A10488) with 5% charcoal-stripped serum (CSS, Gemini, #100-119), twice a day for three days. For genome-edited models with multiple clones, individual clones with the same genotypes were equally pooled for subsequent experiments.

Generation of BCK4 cells has been previously reported^5^. Cell lines were maintained in the following media (Life Technologies) with 10% FBS: MDA-MB-468 in DMEM, MDAMB-134 and MDA-MB-330 in 1:1 DMEM: L-15, ZR75-1 in RPMI and BCK4 in MEM with nonessential amino acids (Thermo Fisher, #11140050) and insulin (Sigma-Aldrich, #91077C).

**Generation of MCF7 stable overexpression *ESR1* mutation cell model**

To generate *ESR1* mutant overexpression cell models, *ESR1* WT and mutant plasmids in pcDNA3.1 backbone were obtained from Addgene (*ESR1*-HA-WT #49498; *ESR1*-HA-Y537S #49499; *ESR1*-HA-D538G #49500, Empty vector #V790-20). MCF7 parental cells (ATCC, HTB-22) maintained in 10% FBS DMEM were transfected with each of the plasmid and subjected to 500 μg/ml G418 (Thermo Fisher, #10131035) selection for 3 weeks. G418-containing medium were changed every 3 days during the selection process. Overexpression of WT and mutant ER was further examined by immunoblot and ddPCR in pooled clones and used for further experiments.

**Reagents**

siRNA against *ESR1*(L-003401), PGR (L-003433), CTCF (L-020165) and non-targeting scrambled control (D-001820-01) were purchased from Horizon Discovery.

Progesterone (P1030) and RU486 (m8046) were obtained from Sigma-Aldrich.

**Quantitative Real-Time Polymerase Chain Reaction (qRT-PCR)**

Different cell models were seeded into 6-well plate with 120,000 cells per well using biological triplicates. After the respective treatments, RNAs were extracted using Qiagen RNeasy Kit, and cDNAs were synthesized using PrimeScript RT Master Mix (Takara Bio, #RR036). qRT-PCR reactions were performed with SybrGreen Supermix (BioRad, #1726275), and the ΔΔCt method was used to analyze relative mRNA fold changes and RPLP0 levels were measured as the internal control. All primer sequences are shown in Supplementary Table S9.

**Immunoblotting**

Cells were lysed with RIPA buffer plus protease and phosphatase cocktail (Thermo Scientific, #78442) and sonicated. Protein concentration of each sample was determined by Pierce BCA assay kit (ThermoFisher, #23225). 40 ug (ER, HA and PR) or 120 ug (CK5 and CK16) proteins were loaded onto 10% SDS-PAGE gel, and then transferred onto PVDF (ER, HA and PR blot) or NC (CK5 and CK16 blot) membrane. The blots were immune-stained with corresponding antibodies. For ER, HA and PR blots visualization, Licor blot fluorescence scanner was used following IRDye 680LT or 800CW secondary antibodies incubation. For CK5 and CK16 blot, chemiluminescent approach was used following Amersham HRP-linked secondary antibody (Millipore Sigma (GENA934) and Clarity Western ECL substrate (BioRad, #1705061) incubation. Antibodies against ER (#8644), HA-tag (#3724) and PR (#3176) were purchased from Cell Signaling. Tubulin antibody was obtained from Sigma (T6557). Antibody against CK5 (ab52635) and CK16 (ab76416) were from Abcam.

**Immunofluorescent Staining**

MCF7 cells were hormone deprived and seeded on coverslips. After desired treatments, cells were fixed with 4% paraformaldehyde and blocked with 3% BSA solution plus 0.1% tritonX-100. Primary antibody against CK8 (Abcam, ab53280), CK5 (Abcam, ab52635), CK16 (Abcam, ab76416) and CK17 (Cell Signaling Technology, #4543) was applied to stain the cells. For counterstaining, CK16 (Santa Cruz, #53255) and ER (Licor, 6F11) mouse monoclonal antibodies were used to combine with above-mentioned rabbit CK5/16/17 antibodies. Secondary Alexa Fluor 488 or 546-conjugated antibodies (Thermo Scientific, A16079 & A11018) and Hoechst (Thermo Scientific, #62249) were used following primary antibody incubation. Coverslips were mounted and images were taken using fluorescence microscope (Olympus, CZX16) under objective of 20X. CK5/16/17 positivity quantification was performed by dividing cells with full cytoskeleton CK expression to total cell numbers of each image. For ER counterstaining quantification, ER signal intensity was quantified using ImageJ for each CK positive cell and five proximal CK negative cells.

**Chromatin-immunoprecipitation (ChIP)**

ChIP were performed as previously described ^6^. Briefly, hormone-deprived MCF7 WT and mutant cells were treated with vehicle or 1nM E2 for 45 minutes. Chromatin DNA was then extracted from each sample. The immunoprecipitation was performed using CTCF (EMD Millipore, 07-729), ERα (Santa Cruz Biotechnologies, sc543) and rabbit IgG (Santa Cruz Biotechnologies, sc2027) antibodies. For CTCF ChIP, qPCR was employed and fold enrichment method was used to quantify the binding enrichment at the selected sites. All primers used was recorded in Supplementary Table S9. For ER ChIP-seq, pooled DNA samples from individual clones were sent to McGill sequencing core using Illumina Hiseq 2000 Platform.

**Chromatin-immunoprecipitation (ChIP) and sequencing analysis**

ChIP was performed as previously described ^6^. ChIP-seq reads were aligned to either hg38 genome assembly using Bowtie 2.0 ^7^, and peaks were called using MACS2.0 with p value below 10E-5 ^8^. We used Diffbind package ^9^ to perform principle component analysis, identify differentially expressed binding sites and analyze intersection ratios with other data sets. Briefly, all the BED files for each cell line were merged and binding intensity was estimated at each site based on the normalized read counts in the BAM files. The pairwise comparison between WT and mutant samples were performed to calculate the fold change (FC) and the binding sites were sub-classified into three categories: gained sites (FC>2), lost sites (FC<-2), and not-changed sites

(-0.2<FC<0.2).

**Single-cell RNA-sequencing analysis**

Two bilateral bone metastases (BoMs) were collected from a patient initially diagnosed with ER+ primary breast cancer, and immediately dissociated into single cells using tumor dissociation kit from Miltenyi Biotech (130095929) following manufacturer’s protocol. Red blood cell lysis (Qiagen158904) and dead cell removal (Miltenyi Biotech 130090101) were performed according to the manual.

Raw counts were mapped to human genome assembly (version GRCh38) using cellranger count function, and the mapped count matrix was imported into Seurat (v 3.1.4) for further analysis. Genes with detected expression in less than 20 cells, as well as cells expressing less than 300 genes or more than 8,000 genes, or containing more than 45% mitochondrial genes were removed, resulting in 10,056 cells for downstream analysis. Mitochondrial genes were regressed out before principle component analysis, and a shared nearest neighbor optimization based clustering method was used for identifying cell clusters. Cell type of each cluster was assigned by the expression of canonical cell markers, and cell signatures derived from single cells RNA sequencing data of defined cell types collected in PanglaoDB database. Log normalized counts values of S100A8, S100A9, TLR4 and AGER were compared between different cell types.
